# Ramp-to-Threshold Dynamics in a Hindbrain Population Controls the Timing of Spontaneous Saccades

**DOI:** 10.1101/430645

**Authors:** Alexandro D. Ramirez, Emre R.F. Aksay

## Abstract

Organisms have the capacity to make decisions based solely on internal drives. However, it is unclear how neural circuits form decisions in the absence of sensory stimuli. Here we provide a comprehensive map of the activity patterns underlying the generation of saccades made in the absence of visual stimuli. We performed calcium imaging in the larval zebrafish to discover a range of responses surrounding spontaneous saccades, from cells that displayed tonic discharge only during fixations to neurons whose activity rose in advance of saccades by multiple seconds. We lesioned cells in these populations and found that ablation of neurons with pre-saccadic rise delayed saccade initiation. We analyzed spontaneous saccade initiation using a ramp-to-threshold model and were able to predict the times of upcoming saccades using pre-saccadic activity. These findings suggest that ramping of neuronal activity to a bound is a critical component of self-initiated saccadic movements.

## Introduction

While much is known about the manner in which motor systems execute behaviors, relatively little is understood about the neural systems that help us decide upon and plan for particular actions, especially those actions that are spontaneous and self-initiated. Classical work in humans identified the presence of pre-movement signals in EEG recordings that slowly built-up for over a second in advance of a self-initiated action (Kornhuber & Deecke, 1965). Work in animal models suggests that this macroscopic activity is reflective of neuronal firing patterns where rates increase ahead of spontaneous movement in a ramp-like manner (Maimon & Assad, 2006; Okano & Tanji, 1987; Schultz & Romo, 1992a). Furthermore, such studies have shown trial-to-trial variation in the timing of the movement is associated with variation in the ramp rate, with a slower rise coupled to longer rise time, suggesting that action is initiated when neural activity ramps to a cell-specific threshold (Maimon & Assad, 2006; Murakami, Vicente, Costa, & Mainen, 2014). More generally, such ramp-to-threshold dynamics have also been found ahead of goal-directed movements (Lin et al., 2020; Schultz & Romo, 1992b) and in decision-making tasks (Bahl & Engert, 2020; Cain, Barreiro, Shadlen, & Shea-Brown, 2013; Dragomir, Štih, & Portugues, 2020; Gold & Shadlen, 2007; Lin et al., 2020; Scott et al., 2017) where the animal is responding expressly to some externally provided stimulus, leading to the proposal that there may be common principles underlying the generation of ramp-to-threshold activity associated with motor planning for both spontaneous and externally-driven actions (Murakami & Mainen, 2015; Schurger, Sitt, & Dehaene, 2012). However, there are many basic questions unanswered about this ramp-to-threshold dynamic, especially in the spontaneous setting, not only regarding how it is generated, but whether it is preparatory, possibly facilitatory (but not essential), or perhaps suppressive (Ames & Churchland, 2019; Lara, Elsayed, Zimnik, Cunningham, & Churchland, 2018; Wise, Weinrich, & Mauritz, 1986; Wong, Haith, & Krakauer, 2015).

The potential role of ramp-to-threshold dynamics in action initiation has been of particular interest in the visuo-motor system. In this setting such dynamics have been most often investigated in the context of a goal-directed action, the performance of a saccade after a cue is provided to a visual target. Of note, neurons in the lateral intraparietal cortex, frontal eye fields, superior colliculus (or optic tectum), and the pons show, in addition to a burst of action potential at the time of goal-directed saccade, a buildup in firing characteristic of ramp-to-threshold dynamics (Munoz, Dorris, Pare, & Everling, 2000; Watanabe & Munoz, 2013). Lesions in midbrain and forebrain regions lead to delays in goal-directed saccade initiation, consistent with a role for pre-saccadic activity in planning and preparation for saccades (Hikosaka & Wurtz, 1985; Schiller, True, & Conway, 1980). However, it is unclear what the role of these dynamics might be in the generation of spontaneous, self-initiated saccades. Although ramping activity is present on cortical, collicular, and tectal neurons ahead of spontaneous saccades, removal of these regions does not alter spontaneous saccade performance and apparently timing in the dark (Albano, Mishkin, Westbrook, & Wurtz, 1982; Johnstone & Mark, 1969; Tusa, Zee, & Herdman, 1986). This suggests that structures in the hindbrain could be involved in the initiation of spontaneous saccades, an obvious choice being the pons, but potentially as well the cerebellum, where recently neurons associated with self-timing of saccades to visual targets have been found (Ashmore & Sommer, 2013).

To better understand the neural systems underlying the initiation of spontaneous saccades, we turned to a model genetic vertebrate, the zebrafish. Even in the larval state this animal makes robust spontaneous saccades in the dark without training or priming (Easter Jr. & Nicola, 1997). In the larval and juvenile stages the entire brain of the intact animal is optically accessible to advanced imaging modalities, enabling tracking of neuronal firing at sub-cellular resolution and at global scale (Ahrens, Orger, Robson, Li, & Keller, 2013; Dunn, Gebhardt, et al., 2016; Portugues, Feierstein, Engert, & Orger, 2014). Coupled with this optical access, the relatively small scale of the brain facilitates structure-function analyses with single-cell anatomical tracing (Bianco et al., 2012; Lee, Arrenberg, & Aksay, 2015) and electron-microscopy based connectomics (Hildebrand et al., 2017; Vishwanathan et al., 2017). Optical access also enables perturbative strategies like holographic optophysiology (dal Maschio, Donovan, Helmbrecht, & Baier, 2017) and targeted ablations (Liu & Fetcho, 1999; Miri, Daie, Burdine, Aksay, & Tank, 2011; Vladimirov et al., 2018) that examine causality. These advantages are particularly relevant given the commonalities in visuo-motor anatomy and physiology between zebrafish and other vertebrates (Straka & Baker, 2013; Straka & Dieringer, 2004).

Our combined imaging and perturbation study provides three notable findings. One, we generated a comprehensive map of neuronal activity patterns underlying spontaneous saccades and subsequent fixations. Two, we found neurons in the hindbrain whose activity rises above baseline in a direction-selective manner multiple seconds *before* the decision to saccade. Furthermore, the time and rate of rise of these cells is consistent with a ramp-to-threshold model which can be used to predict saccade timing. Third, we found, through targeted, single-cell ablations, evidence implicating these cells in controlling the proper patterning of spontaneous saccades. These findings not only provide insights into the mechanisms controlling the choice of when to shift gaze, but also establish a new model system for understanding the neuronal processes underlying spontaneous, self-initiated actions.

## Results

In the first part of the results we present an analysis of spontaneous eye movements in the dark in larval zebrafish (Fig. 1) and a map of the temporal dynamics of neurons that were active during this behavior (Fig. 2–3). We then analyze the temporal properties and spatial distribution of a special class of neurons found in this map whose activity suggests they play a role in eye movement initiation (Fig. 4–6). We end by systematically mapping how eye movement patterning is affected by ablations of these neurons (Fig. 7).

**Figure 1.**
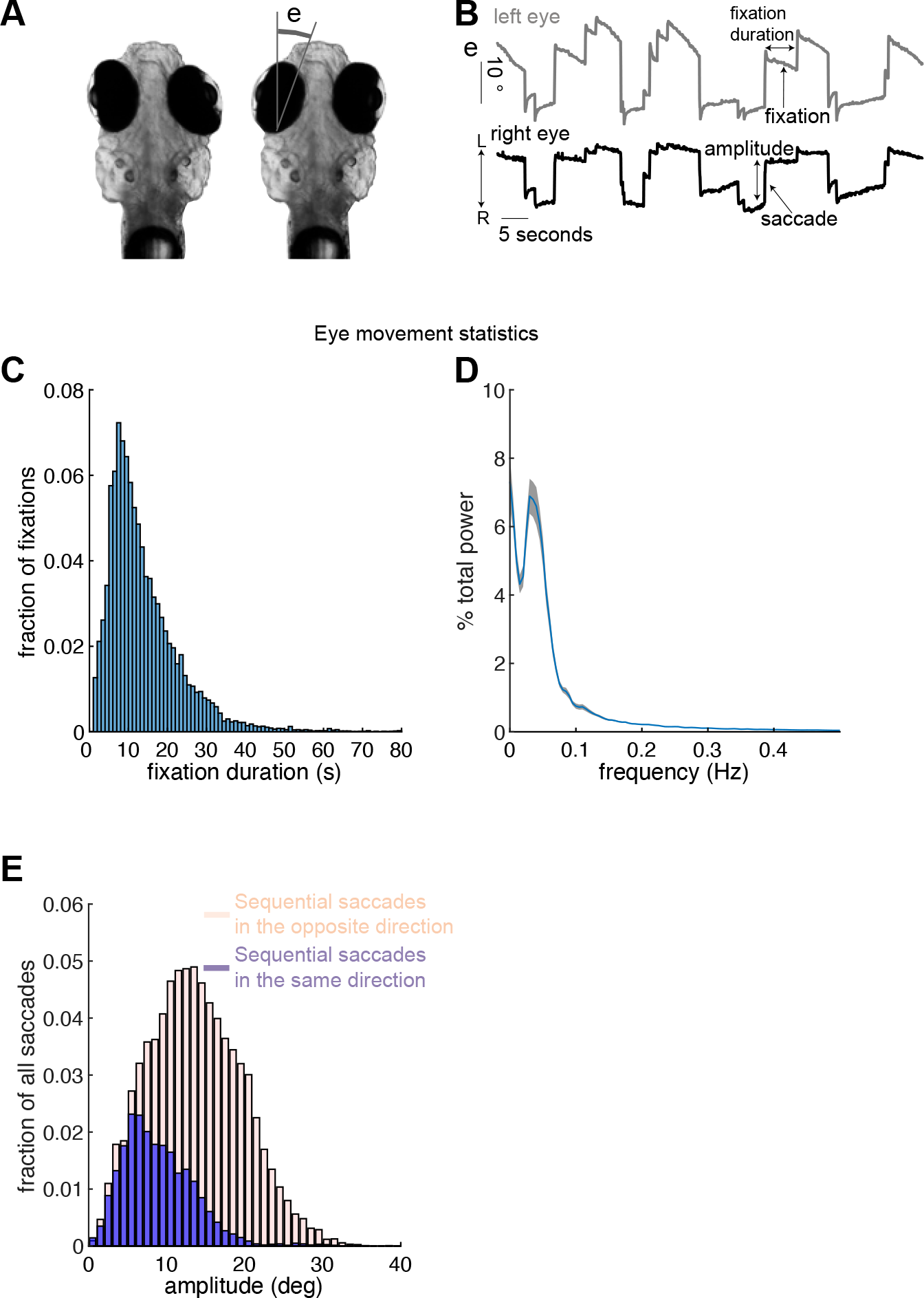
Spontaneous eye movements in larval zebrafish. (A) Changes in horizontal eye position (e) were recorded from agarose-restrained larval zebrafish while they made spontaneous eye movements in the dark. (B) Example changes in horizontal eye position with time for both eyes. Eye movements were quantified by examining saccade amplitude and fixation duration. Leftward/rightward directed saccades are made toward the direction labelled L/R. (C) Histogram of fixation durations measured across multiple animals (1 second bins). (D) Power spectral density of eye-position changes as a function of frequency averaged across all fish and recording sessions (blue; n=422 sessions). Grey shading shows standard error about the mean. (E) Saccade amplitude distribution measured in degrees (1 degree bins). We have separated saccades made in the same direction as the previous saccade (blue) from those made in the opposite direction (beige). Both histograms were normalized by the total number of saccades (n=16,033 saccades).

### Larval zebrafish generate spontaneous eye movements with a range of fixation durations

Larval zebrafish reliably generated spontaneous eye movements consisting largely of a back-and-forth alternation of rapid eye movements known as saccades followed by longer periods of relatively constant eye position known as fixations (Fig. 1A, 1B). Spontaneous eye movements in the horizontal plane were measured in agar mounted larval zebrafish (n=20 fish; 7-8 days post fertilization) after releasing the eyes from the agarose (Fig. 1A). Experiments were performed in the dark to ensure that movements were self-initiated and not occurring in response to a visual target. The time between saccades, known as the fixation duration (see Fig. 1B), was variable; fixations lasted between 2-50 seconds (1 and 99% quantiles) with a median duration of 11.4 seconds (n=16,033 fixations; Fig. 1C). While there was a clear alternating cycle of leftward then rightward directed saccades (Fig. 1B), we found that it was not uncommon for eye movements to deviate from this simple pattern, as nearly a quarter of the time zebrafish made successive saccades in the same direction (22.9 + −0.1% average + −sem across fish). To further examine the rhythmicity of these movements, we performed a Fourier analysis of the changes in eye position with time and found that power was distributed over a range of frequencies, with 95% of total power between 0-0.35 Hz and peak power (excluding 0 Hz) at 0.03 Hz (Fig. 1D). The total range of eye position angles was 20 + −1 degrees (average + −sem across fish of the 1^st^ and 99^th^ position quantiles averaged across both eyes). The size of saccades, known as the saccade amplitude (see Fig. 1B), varied between 2-28 degrees (1 and 99% quantiles, median amplitude = 12 degrees) and depended on the direction of the previous saccade (Fig. 1E; median amplitude of saccade in the same direction as previous saccade equaled 8 degrees; median amplitude of saccade in the opposite direction equaled 13 degrees). These results show that larval zebrafish can self-initiate a simple yet varying pattern of horizontal eye movements.

### Mapping Activity Using Two-Photon Microscopy

To create a comprehensive map of the functional cell types involved in self-initiated eye movements, we imaged calcium dynamics throughout the larval zebrafish hindbrain (Fig. 2Ai, 2Aii) during spontaneously generated saccades and fixations in the dark. The seconds-long fixation durations between eye movements (Fig. 1B, 1C) facilitated the use of calcium imaging to analyze changes in single-cell activity during fixations. Activity from single neurons expressing nuclear-localized GCaMP6f under the control of the HuC pan-neuronal promoter was measured with raster-scanning two-photon microscopy (Fig. 2Aiii). In each fish, we imaged a portion of the hindbrain using a stack of 5-68 horizontal planes. Stacks from individual fish were then registered to a reference brain (Supplemental Fig. 1A, see Methods). The planes were centered at different rostral-caudal locations across fish (imaging 20 fish in total) so that when combined via registration we effectively sampled responses from the entire hindbrain (Supplemental Fig. 1B), with each voxel sampled from at least 3 fish (Supplemental Fig. 1C)

**Figure 2.**
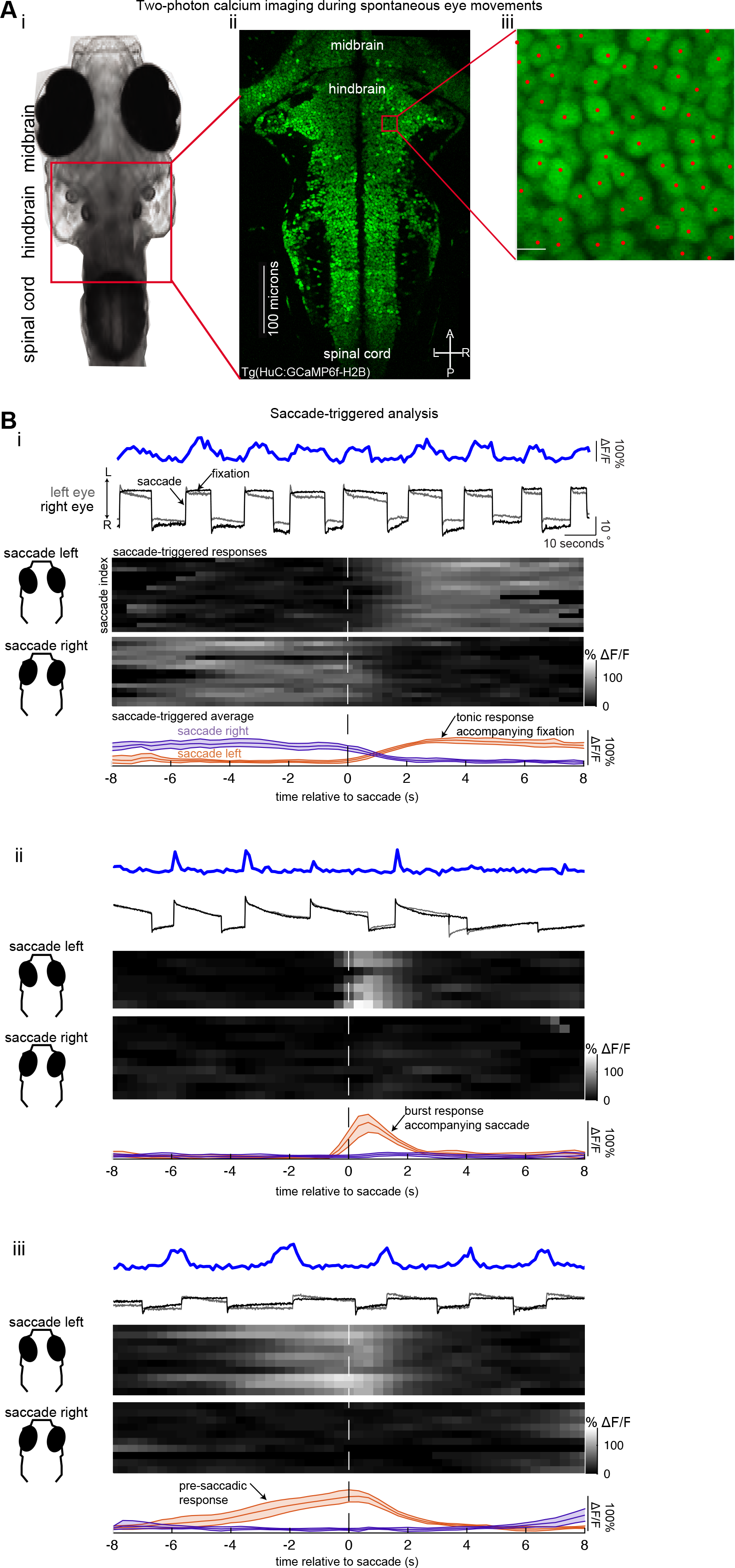
Two-photon calcium imaging during spontaneous eye movements. (A)(i) Brightfield image of a 7-day old larval zebrafish. The location of the hindbrain is outlined by the red box. (ii) Time-averaged calcium image showing all neurons within a single plane of the hindbrain. A, P, L, R denote anterior, posterior, left and right respectively. (iii) Zoomed-in image of time-averaged nuclear-localized GCaMP6f expression showing typical resolution used. Red dots show the center-of-mass for automatically detected nuclei using a morphological opening based computer vision algorithm (see methods). Scale bar length equals 5 microns. (B) Examples of simultaneously recorded eye movements and calcium activity along with saccade-triggered responses and averages. (i-iii, top row) Simultaneously recorded left and right eye position (gray and black) and single-cell fluorescence traces (blue) versus time. Leftward/rightward directed saccades are made toward the direction labelled L/R. (i-iii, middle row) Fluorescence trace is segregated into responses that occur before and after each saccade. Heatmap of saccade-triggered responses around saccades to the left (upper middle) and saccades to the right (lower middle). Grayscale color in each row displays dF/F as a function of time around saccade. White dashed line shows time of saccade. Temporal axis shown in the bottom row. (i-iii, bottom row) Saccade-triggered averages around saccades to the left (red) and right (purple) along with 95% confidence intervals. (i) Example of a cell with tonic post-saccadic activity. (ii) Example of a cell that displays a burst of activity triggered during the saccade. (iii) Example of a cell that displays pre-saccadic activity.

We found that approximately a quarter of the neurons in the hindbrain showed some activity under these conditions. We used standard computer vision algorithms that analyzed time-averaged fluorescence intensity maps (see Methods) to count the total number of cells sampled (Fig. 2Aiii and Supplemental Fig. 1D show cells detected via this method). We measured 243,459 hindbrain cells from all planes and fish in our data set. This number is larger than the 120,000 neurons expected from sampling 3 complete hindbrains because some regions were sampled more than 3 times (Supplemental Fig. 1C). To infer how many of the total cells were active, we performed a separate analysis using an algorithm that relies on non-negative matrix factorization (NMF) to identify the locations of responsive neurons (Pnevmatikakis et al., 2016). As expected, we found that fluorescence activity had fewer deviations on average from cells identified by the computer vision methods that were labelled as background by the NMF algorithm (Supplemental Fig. 1D, 1E). Considering these background neurons as inactive, we determined that approximately a quarter (63,644 hindbrain cells) of hindbrain neurons in our data set were spontaneously active. In the rest of the manuscript, we ignore inactive cells and analyze cells identified by the NMF algorithm (see Supplemental Fig. 1F for a workflow of the cells analyzed in the manuscript).

### Neuronal Activity Dynamics Associated with Spontaneous Saccades and Fixations

Approximately ten percent (n=6,900) of the spontaneously active hindbrain neurons had average activity related to eye movements. We aligned fluorescence responses from active cells to the times of spontaneous saccades and determined which neurons had significant saccade-triggered average (STA) activity. We treated individual fixations surrounding a saccade as single-trials (Fig. 2Bi–iii, Methods) and performed a one-way ANOVA to compare the STA at different time bins to search for cells with significant deviations from baseline. Significant STAs had clear changes in dF/F around or at the time of saccade (Fig. 2Bi–iii). We will refer to cells with significant STAs as eye-movement responsive.

Eye-movement responsive cells had activity patterns that ranged from tonic activity during the post-saccadic epoch (Fig. 2Bi), to burst activity only at the time of saccade (Fig. 2Bii), to activity that gradually climbed during the pre-saccadic epoch (Fig. 2Biii). We used Principal Component Analysis (PCA) to help represent how these activity patterns were distributed across the population. We found that 92% of the variance in STAs could be explained by three components (Fig. 3A). Therefore, we examined the distribution of coefficients along these components (Fig. 3B). We normalized the coefficients so that all responses had the same unit magnitude (Fig. 3Bi-ii) in order to focus on variations in profile shapes (see Methods). The coefficients for the population were broadly, and mostly continuously, distributed (Fig. 3Biii-iv).

**Figure 3.**
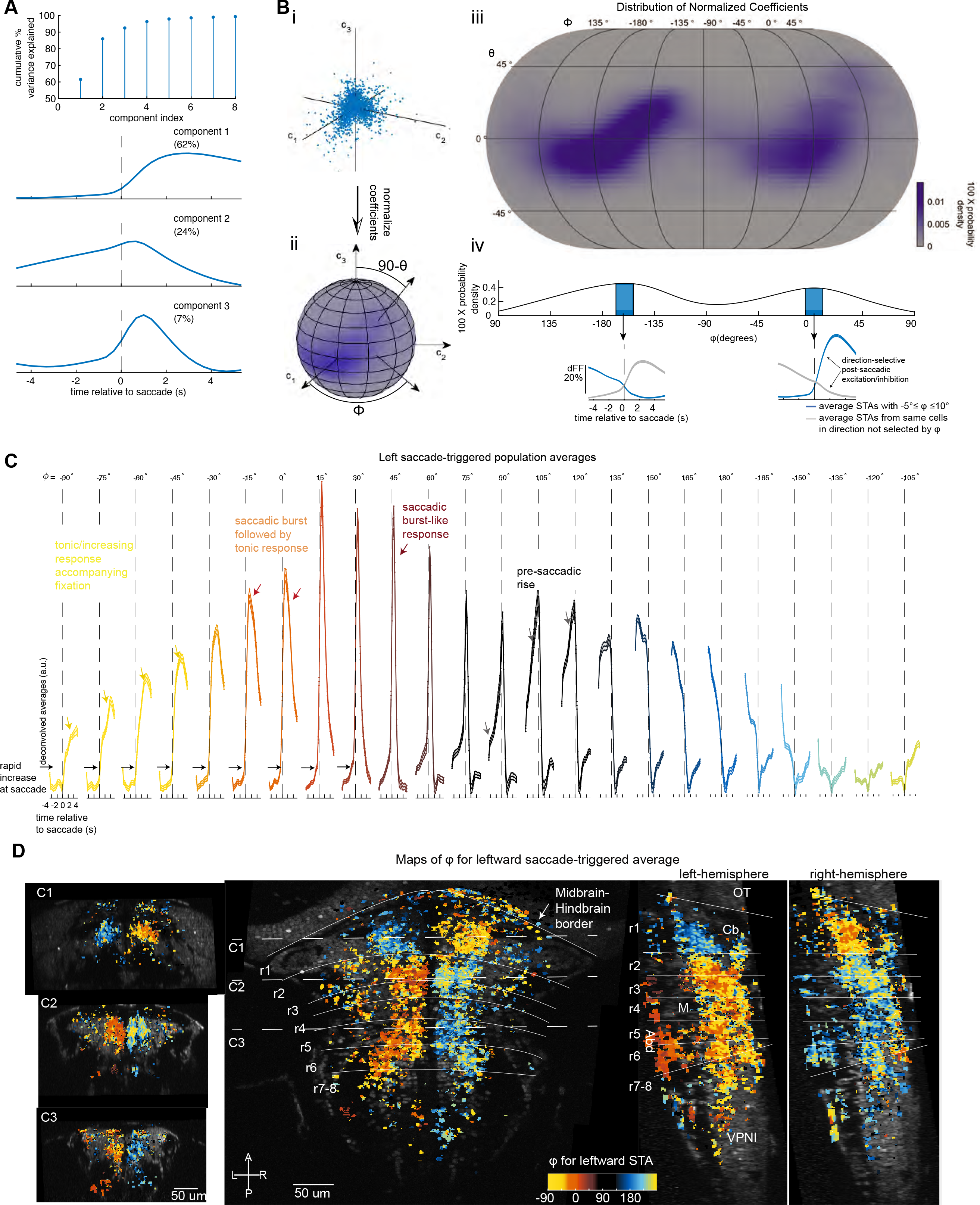
Functional and spatial distribution of neuronal responses. (A) (top row) Cumulative percent variance explained by each component found via a principal component analysis on saccade-triggered averages (STAs). (bottom rows) First three components sorted according to the amount of variance each component explains from most (component 1) to least (component 3). Percentages reported in the labels in the upper right denote the amount of variance each component explains. (B) (i) Scatter plot of coefficients obtained from projecting STAs along components one (c_1_), two (c_2_), and three (c_3_). Each dot represents a different STA. Note that we construct two STAs for each cell, one around saccades to the left and one around saccades to the right. Each cell therefore contributes two dots to the plot. (ii) The length of each coefficient vector was normalized to one which allows each STA to be described as a point on a sphere specified by longitude and latitude, *φ* and *ϑ* respectively. (iii) Two-dimensional probability density of the distribution of normalized coefficients shown on a map using an Eckert IV projection. The angles are chosen so that an STA whose temporal profile is completely characterized by component one, two, or three would have coefficients equal to (*ϕ* = 0, *ϑ* = 0), (*ϕ* = 90, *ϑ* = 0), (*ϕ* = 0, *ϑ* = 90) respectively. (iv) One-dimensional probability density of longitudinal angle, *φ*. The population average STAs for responses corresponding to peaks in the density plot of longitudinal angles (*ϕ* = 3 and −162 degrees) are shown in blue below the density function (individual STAs used to construct each average have values of *ϕ* within 15 degrees, shaded blue region, of the most probable values). Note that there are no restrictions on whether the STAs averaged together to form the blue trace are around saccades to the left or right. Since single-cells have direction-selective responses (see Supplemental Fig. 2 for further analysis), the two STAs for a given cell will have different temporal profiles (one ON and one OFF) each characterized by different coefficient values and thus a single-cell will only contribute one out of its two STAs to the average. For completeness, we show in gray the population average STA that results when grouping together and then averaging the STAs not included in the blue trace using the same cells. (C) Series of population-averaged STAs after deconvolving with an exponential kernel. Population averages are comprised of STAs around saccades to the left specified by the value of *ϕ* shown in the top row, (plus or minus 7.5 degrees). The filled region shows one standard error about the population average. Responses are color coded to enable evaluation of spatial organization in D. (D) Horizontal, sagittal and caudal projections of cell locations color coded by the average value of *ϕ* quantifying STAs around saccades to the left. Color scheme corresponds to the colors shown in C. Coronal sections show average values of *ϕ* for cells within 30 microns of the dashed lines marked C1, C2, and C3 in the horizontal map. Sagittal projections show average values of *ϕ* quantifying STAs around saccades to the left for cells in the left hemisphere or right hemisphere. Number of fish in the averages depends on cell location (see Supplemental Fig. 1C). r, rhombomere; OT, optic tectum; Cb, cerebellum; M, Mauthner cell; VPNI, velocity-to-position neural integrator; Abd, Abducens complex; L-R, left-right; A-P, anterior-posterior.

A clustering analysis of the coefficients found that these data were best segregated into two clusters corresponding to left and right selectivity. The peaks in the density of coefficients corresponded to two distinctive STAs; one STA increased in dF/F following saccades while the other STA decreased (Fig. 3Biv, blue traces). To measure how these patterns in STAs related to patterns in single-cell responses, we performed a K-means analysis on the combined coefficients for the two STAs, those around saccades to the left and right, that characterize single-cell responses. We found evidence that the coefficients for single cells were best segregated by two clusters (Supplemental Fig. 2A, 2B). The cluster means revealed that a typical cell has an STA that increases around saccades in one direction (left or right) and decreases around saccades in the other direction (Supplemental Fig. 2C). This direction-selective response captured the responses seen in most cells (see the cluster separation in Supplemental Fig. 2C and quantified in Supplemental Fig. 2B), indicating that the peaks in the density of coefficients (Fig. 3Biii,iv) result from neurons that selectively respond to either saccades to the left or saccades to the right.

We next generated population averages of STAs grouped according to their coefficient, *φ*, see Fig. 3Bii and Methods section *Principal Component* Analysis, to ease visualization of both the dynamics across the population and the spatial distribution of the different activity profiles. STAs around saccades to the left were ordered by *φ*, grouped into bins of 15 degree width, and then averaged together and plotted in Figure 3C. The equivalent plot for saccades to the right is shown in Supplemental Figure 3A (in both plots, each bin contained at least 1% of eyemovement responsive cells - the qualitative trends described below remained largely unaffected by sub-dividing groups further, see Supplemental Fig. 4). Before population averaging, we deconvolved the STAs of these cells using an exponential kernel with two-second time-constant to approximate the effects of GCaMP6f calcium filtering on their average firing rate (Kawashima, Zwart, Yang, Mensh, & Ahrens, 2016; Yaksi & Friedrich, 2006). We also registered the cell locations to a zebrafish brain atlas (Randlett et al., 2015) to map cell dynamics to neuronal locations (Methods).

We found stereotyped patterns of eye-movement related activity at specific spatial locations. Many cells had pre-saccadic activity that remained constant at baseline levels, increased within a second of the saccade, and then displayed tonic firing after the saccade. Some of these neurons displayed primarily ‘tonic’ firing after the saccade (e.g. *φ* = −75 in Fig. 3C and Supplemental Fig. 3A); others had more ‘burst-tonic’ characteristics (e.g. *φ* = 0). Both cell types were broadly distributed in the hindbrain (Fig. 3D, Supplemental Fig. 3B), with tonic neurons generally located at more dorsal locations than burst-tonic neurons. Prominent pockets of tonic and burst-tonic neurons with ipsilateral sensitivity were found in rhombomeres 7 and 8 in regions previously associated with the velocity-to-position neural integator (VPNI in Fig. 3D), ventral portions of rhombomeres 5 and 6 associated with the abducens complex (ABD in Fig. 3D), dorsal regions of rhombomeres 5 and 6, and dorsal regions of rhombomeres 2 and 3; neurons with contraversive tonic or burst-tonic responses were located in rhombomere 1 below the cerebellum. Other neurons had purely ‘burst’ activity associated with ipsiversive saccades as expected for saccade generator cells (*φ* = 45 in Fig. 3C and Supplemental Fig. 3A); these were most prominently clustered in ventral portions of rhombomeres 2 and 3. With increasing *φ*, we observed a pre-saccadic rise in activity that becomes longer in duration and more prominent relative to the burst event (*φ* = 105 in Fig. 3C and Supplemental Fig. 3A). This pre-saccadic rise is suggestive of activity that is involved in the timing of upcoming saccades, a role which we will explore further in the remainder of this paper.

In summary, we found that about 3% of hindbrain neurons in larval zebrafish had responses associated with spontaneous saccades and fixations, these responses were direction-selective, and the response profile of this population was diverse. The diversity included cells with step-like profiles expected for abducens neurons and integrator neurons, and cells with burst-like responses expected for saccade generator neurons. The distribution also contained neurons whose activity is better described as anticipating upcoming movements, consistent with a role in motor preparation, a role which we explore below.

### Single-Trial Analysis of Cells with Pre-Saccadic Rise Activity

Within the continuum described above were cells whose population-average activity steadily rose ahead of the upcoming saccade. Because this form of dynamics appears to anticipate a future movement, it is reasonable to speculate that neurons with such activity play an important role in saccadic preparation or timing. We now turn our attention to a closer examination of such neurons, with a focus on single-trial and single-cell level analyses that can capture variations missed with averaging.

We found that 6% (n=430) of eye-movement responsive hindbrain neurons had fluorescent activity with clear ramp-like activity ahead of saccades (Fig 4A). To systematically select these cells, we re-analyzed eye-movement responsive neurons (see workflow in Supplemental Fig. 1F) whose Spearmen correlation coefficient between fluorescence activity and time to upcoming saccade was significantly greater than zero (p<0.01 using the Holm-Bonferroni method to correct for multiple comparisons). Here we note some of the key qualitative features of these neurons before providing detailed quantification. These cells showed clear increases in activity before individual saccades either to the left (Fig. 4A, neurons in 3^rd^ and 4^th^ row from top) or right (Fig. 4A, neurons in 1^st^ and 2^nd^ row from top). For each cell, pre-saccadic rise events occurred ahead of saccades in one “ON” direction, although there were occasions where cell activity also rose before saccades in the opposite direction or failed to rise before the ON direction. dF/F for a small number of cells was significantly correlated with upcoming saccades in both directions (n=4 cells, not shown). Rise events generally lasted multiple seconds, and there was variability in the duration and rate of the rise both within and across cells. Further, for a given cell, activity tended to rise to consistent values at the time of saccade, even though duration of rise varied by 50% or more. We will refer to these cells as pre-saccadic rise (SR) neurons.

**Figure 4.**
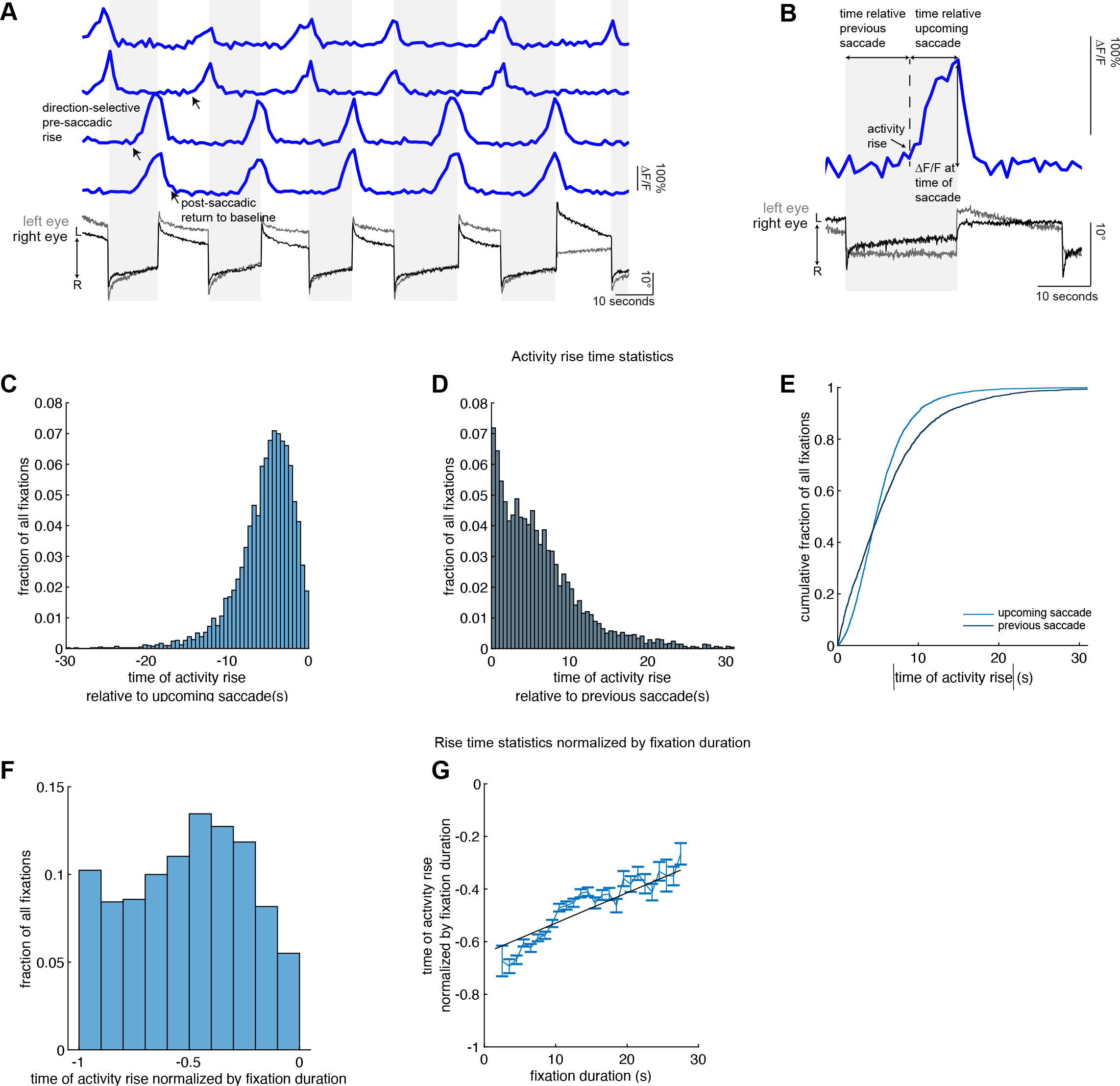
Neurons whose activity anticipates saccades in a direction-selective manner. (A) Simultaneously recorded left and right eye position (gray and black) and single-cell fluorescence traces (blue) versus time from cells whose dF/F values rise before spontaneous saccades. In panels A and B, leftward/rightward directed saccades are made toward the direction labelled L/R. (B) Zoomed-in view of pre-saccadic fluorescence activity demonstrating the measurements of spontaneous pre-saccadic activity and relevant features used in subsequent analyses. (C) Histogram of times of pre-saccadic activity rise relative to the time of upcoming saccade. (D) Histogram of times of pre-saccadic activity rise relative to the time of the previous saccade. Bin size in C and D is 0.5 seconds. (E) Comparison of the cumulative distribution functions of times of pre-saccadic activity rise relative to upcoming or previous saccade. We take the absolute value of time of pre-saccadic activity rise relative to the time of upcoming saccade, which is strictly negative by construction, in order to compare the distributions. (F) Histogram of times of pre-saccadic rise, measured with respect to upcoming saccade, normalized by fixation duration (bin size is 0.1). (G) Average time of pre-saccadic rise normalized by fixation duration as a function of fixation duration (blue line, 1 second bins). Error bars show standard error about the mean. Line of best fit linear regression model shown in black.

To quantitatively characterize the dynamics of SR cells, we assessed several features of their activity before and at the time of saccade that could elucidate their role in initiating the upcoming spontaneous saccade (Fig. 4B, Methods). In order to examine how the initiation of activity related to saccade occurrence, we measured the time when activity rose above baseline, and compared that to the time of upcoming and previous saccades. We also measured whether the time of activity rise scaled with fixation duration. To determine whether neuronal activity consistently rose to similar values, we measured activity at the time of saccade.

The initiation of the rise event was more tightly coupled to the timing of the upcoming saccade rather than the timing of the previous saccade. Across cells and fixations, activity rose over a range of times between 2 and 10 seconds relative to the upcoming saccade (10 and 90% quantiles; median time = 5 seconds, n= 4,161 fixations across 15 fish; distribution shown in Fig. 4C). The timing of activity initiation when measured relative to the occurrence of the previous saccade was generally more variable, with a range of 1 to 13 seconds and with a variance that was three times larger than the variance in timing when measured with respect to upcoming saccade (Fig. 4D). Furthermore, we rejected the null hypothesis that these two measurements in the time of rise come from the same distribution (Fig. 4E, p<0.001; two-sample KS-test). A similar trend was observed when data was analyzed on a per cell basis: 69% of cells had activity whose time of rise was more variable when measured relative to the previous saccade than when measured relative to upcoming saccade (ratios greater than one in Supplemental Fig. 5A). Therefore, the activity of SR neurons is more informative of upcoming saccades than previous saccades.

We found that cell fluorescence did not rise at a fixed time relative to saccades. Given that there was notable variation in the fixation duration (Fig. 1C), we also tested for relationships between the time of rise and the duration of the fixation. To see if there was a constant scaling relationship between fixation duration and the time of SR activity rise, we examined whether the distribution of times of rise normalized by fixation duration was peaked (Fig 4F). We found that, across cells and fixations, normalized rise times were not peaked at a single value but rather distributed across the full range of possible values. The variability in normalized time of rise was related to fixation duration (Fig. 4G): there was a slight trend for activity to rise relatively quickly after the previous saccade (normalized times near −1) during short fixation durations and for activity to rise later in the interval during longer durations (Fig 4G best fit line slope = 0.012 + −0.001(1/s) and offset = −0.645 + −0.008, estimate + −sem). On a per cell basis, time of normalized rise and fixation duration appeared monotonically related for some cells but not others (see Supplemental Fig. 5B; 22% of cells had a Spearman correlation coefficient above 0.5, see Supplemental Fig. 5C). While the distribution of correlation coefficients across cells was significantly different than shuffled controls (Supplemental Fig. 5C), within-cell variability made it difficult to discern significant linear trends (linear regression found 19% of cells had slopes with a t-test p-value <0.05, Supplemental Fig. 5D). In summary, we found that SR cell activity can rise within any fraction of the fixation time, with a propensity, at the population level, to begin rising later in the fixation for longer fixation durations.

### Pre-Saccadic Rise Dynamics are Predictive of Upcoming Saccades

If SR cell activity determines when a spontaneous saccade should occur, we should be able to predict whether a saccade is about to happen based on the output of SR populations. In this section, we show that SR pre-saccadic dynamics can be used to predict saccade direction and time. As part of this analysis we also detail the relationships between the time at which an SR cell begins rising ahead of a saccade and the speed at which activity rises.

We first quantified how well SR cell activity can predict the direction of upcoming saccade. As an initial assessment of the information SR populations contain regarding upcoming saccade direction (left or right), we quantified population fluorescence using the choice probability (Crapse & Basso, 2015; Latimer, Yates, Meister, Huk, & Pillow, 2015). The choice probability predicts saccade direction by comparing SR population average activity before saccades in the preferred direction with population activity before saccades in the non-preferred direction (see Methods). We found that choice probability, averaged across fixations, increased monotonically with time when aligned to saccade onset (Fig. 5A). Near the saccade, the choice probability was well above chance levels of 50% (Fig. 5A; mean choice probability equals 79%). An ideal observer who has knowledge of the saccade transition probabilities would guess that the saccade is in the opposite direction of the previous saccade and be correct 77% of the time (given that successive saccades occur in the same direction 23% of the time, see section “Larval zebrafish generate spontaneous eye movements with a range of inter-saccade times”). Therefore, the choice probability performance is on par with the performance of an ideal observer. These results indicate that spontaneous SR population activity contain information regarding the upcoming saccade direction.

**Figure 5.**
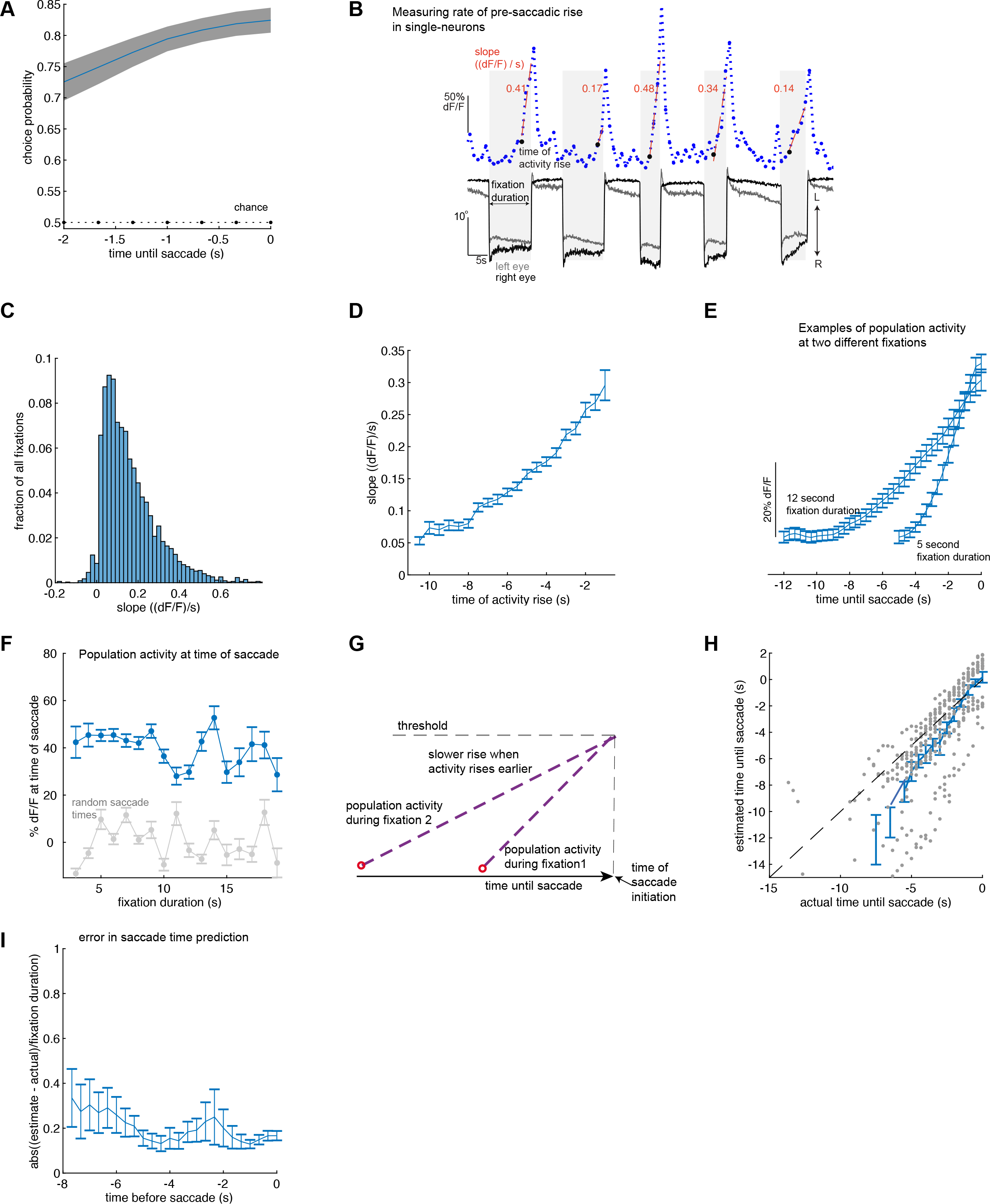
The rate and time of pre-saccadic rise vary in a manner consistent with ramp-to-threshold models. (A) Choice probability averaged across all fixation durations between 2 and 20 seconds as a function of time until upcoming saccade (blue line). Dotted black line shows choice probability from randomly guessing saccade direction. (B) Visualization of the slope statistic (orange) used to quantify rate of pre-saccadic rise before each saccade. Simultaneously recorded fluorescence activity (blue) is plotted along with eye movements. Leftward/rightward directed saccades are made toward the direction labelled L/R. (C) Histogram of rates of pre-saccadic fluorescence increase. Bin size is 0.02 (dF/F)/s. (D) Average slope of pre-saccadic fluorescence increase as a function of time of pre-saccadic activity rise measured with respect to upcoming saccade. Bin size is 500ms. (E) Example traces of population average, pre-saccadic fluorescence as a function of time until upcoming saccade. Fixation duration was held fixed to either 5 seconds (to within 0.5 seconds; n=263 fixations from 119 cells across 10 fish) or 12 seconds (to within 0.5 seconds; n= 177 fixations from 148 cells across 13 fish). (F) Population average fluorescence at the time of saccade as a function of fixation duration (blue; number of fixations in average varies between 43 and 308). Gray trace shows population average fluorescence of the same cells after replacing saccade times with randomly selected times. (G) Hypothetical mechanism for how a pre-saccadic signal could initiate saccades. Activity from a population of neurons (purple dashed lines) initiates saccades once it crosses an intrinsic threshold value (gray dashed line). Activity rises later in fixation 1 than in fixation 2. Since pre-saccadic activity in fixation 2 is assumed to reach the same threshold as pre-saccadic activity in fixation 1, activity in fixation 2 must rise more slowly than activity in fixation 1. (H) Predicted time until saccade versus actual time (gray dots). Average predicted time versus time until saccade shown in blue (500ms bins) (I) The average prediction error of the gray dots in (H) as a function of actual time until saccade. Prediction error is measured relative to the fixation duration. Error bars in A (gray shading), D, E, F, H and I show standard error about the mean.

We next examined the relationship between the time at which an SR cell began ramping and its rate of rise, finding that these dynamics generally fit under a “ramp-to-threshold’ framework. For each cell and fixation, we measured the rate of rise of pre-saccadic activity (Fig. 5B, Methods). Across cells and fixations, the rate of rise varied between −0.03−0.60 (dF/F)/s (1 and 99% quantiles, mean=0.16, n= 3,483 events from 415 cells across 15 fish; distribution shown in Fig. 5C). This variability was positively correlated with the duration of ramping activity: slowly rising activity began to increase above baseline on average earlier than faster rising activity (Fig. 5D and 5E; at the individual cell level, this positive correlation was apparent for 29% of SR cells, Supplemental Fig. 5E, 5F). We also examined SR activity conditioned on whether two consecutive saccades occurred in the same or opposite direction (Supplemental Fig. 6A) and found similar trends in the distribution of the rate of rise (Supplemental Fig 6B) and the correlation between rate of rise and fixation duration (Supplemental Fig. 6C). Furthermore, the average activity reached similar levels at the time of saccade (Fig. 5F). These data suggest a picture where spontaneous saccades are triggered when the entire population of SR activity reaches a threshold, with rapid rates of rise associated with a shorter time to saccade initiation (Fig. 5G).

We next used the ramp-to-threshold framework to quantify how well SR neurons predict upcoming saccade times. We predicted saccade times by calculating when a ramp-to-threshold model, in which population activity growth is approximated as linearly increasing before saccade, passes a constant threshold level associated with saccade initiation (see Methods). We found good agreement between population activity and the ramp-to-threshold model when we trained the model using the entire period when SR activity increases (*r*^2^=0.69 see Supplemental Fig. 7; 40% of cells were held-out for testing; 60% of cells were used to train the ramp rate). We next trained the model using a running estimate of the ramp rate, as one would do to predict saccade times in real-time. We found that the predicted times to saccades and the actual times to saccade were well correlated (Fig. 5H, cc=0.61 (p<0.001)), with a timing error of around 20% (Fig. 5I), even when we restricted our training window to the first two seconds after ramp initiation (cc=0.47 (p<0.001)). By contrast, an ideal observer who has knowledge of the time of previous saccade and knowledge of the fixation duration probability distribution predicted saccade times with a timing error of 51% and could not produce predictions that are correlated to the actual saccade times (cc=0.00; see Methods). Thus, the pre-saccadic dynamics of SR population activity can be used to predict when saccades will occur.

### Spatial Distribution of Neurons with Pre-Saccadic Rise Dynamics

We next examined the spatial distribution of hindbrain cells with pre-saccadic rise dynamics. These neurons were broadly distributed across the rostral-caudal axis, with a small group in the cerebellum and rhombomere 1 (8%), the majority between rhombomeres 2-6 (83%), and a small group in rhombomeres 7-8 (9%) (Fig. 6A). SR neurons were more likely to be near the midline than near the edge of the brain (Fig. 6B). Along the dorsal-ventral axis, SR cells largely resided in a region that is 10-70 microns dorsal of the Mauthner cell (Fig. 6C). None of the SR cells were found in the abducens complex (ABD in Fig. 6D), consistent with previous literature on abducens firing rate properties (Pastor, Torres, Delgado-Garcia, & Baker, 1991). Directional preference of pre-saccadic neurons was largely (58%) contraversive (Supplemental Fig. 8), but we did observe clusters of ipsiversive preferring cells. The rostral-caudal distribution of pre-saccadic neurons depended on the cell’s position along the dorsal-ventral axis; nearly all the pre-saccadic neurons located in the caudal portion of the hindbrain (rhombomeres 7-8) were contained near the Mauthner cell level or below (Fig. 6D).

**Figure 6.**
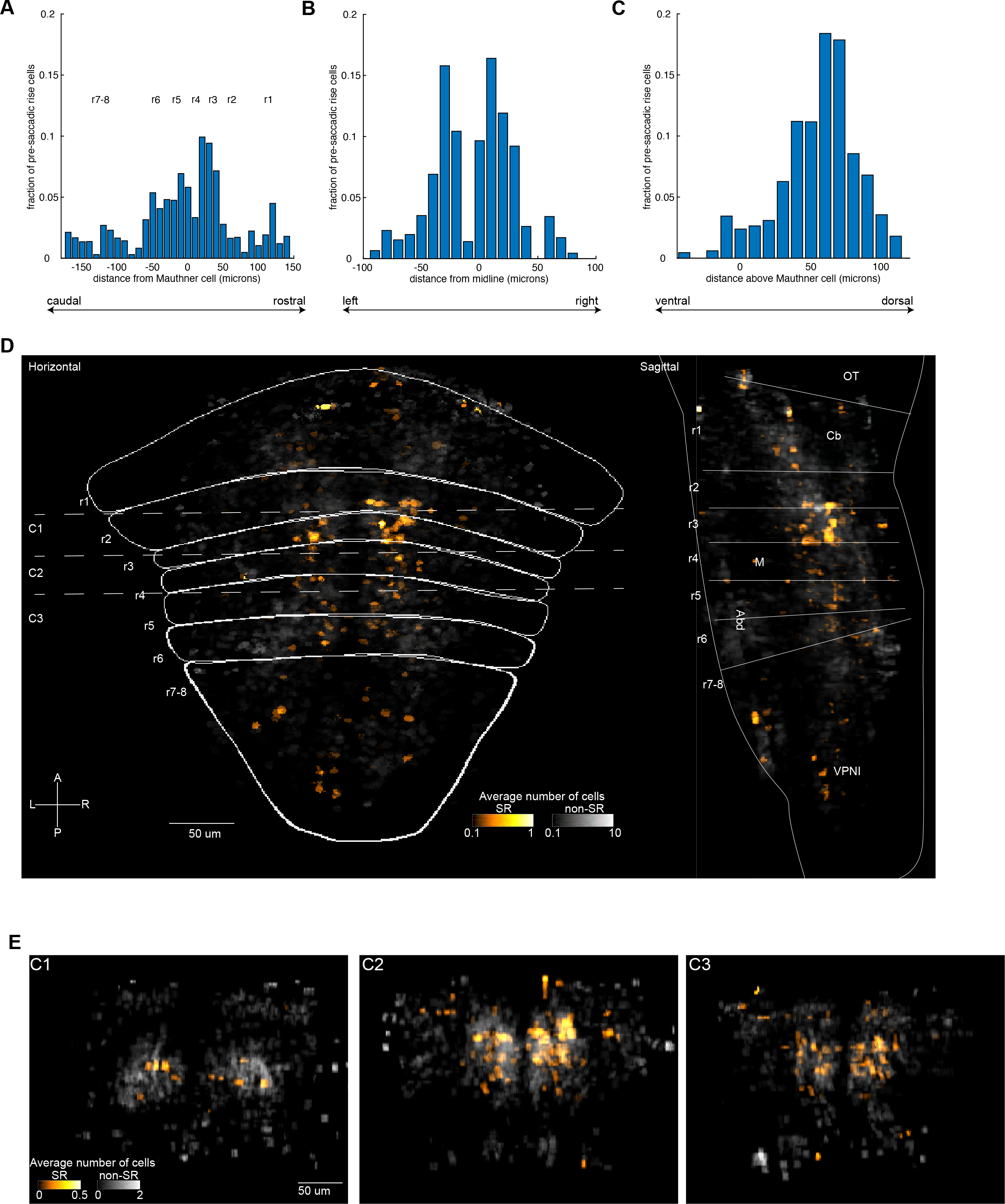
Spatial distribution of cells whose activity anticipates saccades. Histograms showing percentage of pre-saccadic neurons as a function of their registered location onto a common reference brain for the rostral-caudal (A), left-right (B), and dorsal-ventral (C) directions. Percentages are weighted to account for unequal numbers of fish used to sample different hindbrain regions. Bin size is 10 microns for all three panels. (D) Horizontal and sagittal projections showing the average number of SR cells (yellow-orange color scheme) found at each location following alignment with a reference brain. SR projections are on top of horizontal and sagittal projections of the average number of eye-movement responsive cells that are not classified as SR (black-white color scheme). Number of fish in the averages depends on cell location (see Supplemental Fig. 1C). r1-8 indicates approximate rhombomere location within the reference brain. OT, optic tectum; Cb, cerebellum; M, Mauthner cell; VPNI, velocity-to-position neural integrator; Abd, Abducens complex; L-R, left-right; A-P, anterior-posterior. (E) Coronal sections showing SR locations (yellow-orange color scheme) on top of eye-movement responsive, non-SR cells (black-white color scheme) at the same location. C1, C2, C3 show averages for cells within 30 microns of the corresponding dashed lines shown in the horizontal map (D).

### Focal Laser Ablations Identify SR Cells as Indispensable for Setting Spontaneous Fixation Durations

It is unknown where the signals to initiate spontaneous saccade arise. The activity of the SR neurons we identified suggest that they initiate saccades when their population level activity reaches a threshold value. In the simplest instantiation of this model, activity is summed across cells and compared with a threshold value. If true, losses in the number of SR cells would lead to a longer time until saccades occur.

To test this possibility, we performed bilateral two-photon laser ablations at different locations in the hindbrain where SR neurons are found and monitored changes in saccade and fixation metrics. We performed two sets of ablation experiments. In the first, we ablated cells that were clustered together in a volume that was approximately cylindrical in shape with a diameter of 30 micrometers along the rostral-caudal and medial-lateral axis and a side-length of 60 microns along the dorsal-ventral axis (Fig. 7A). Within a single animal, two clusters (one on the left and one on the right hemisphere) were ablated at approximately the same rostro-caudal position (approximately 1,200 cells, or 3% of the hindbrain, ablated per experiment). The rostro-caudal position of the cluster ablations was varied across animals. In the second set of experiments, we targeted single-cells and removed 6-20 individual neurons based on their relationship to eye movements (Fig. 7B). In both sets of experiments, we measured changes in fixation duration following ablations (Fig. 7C).

**Figure 7.**
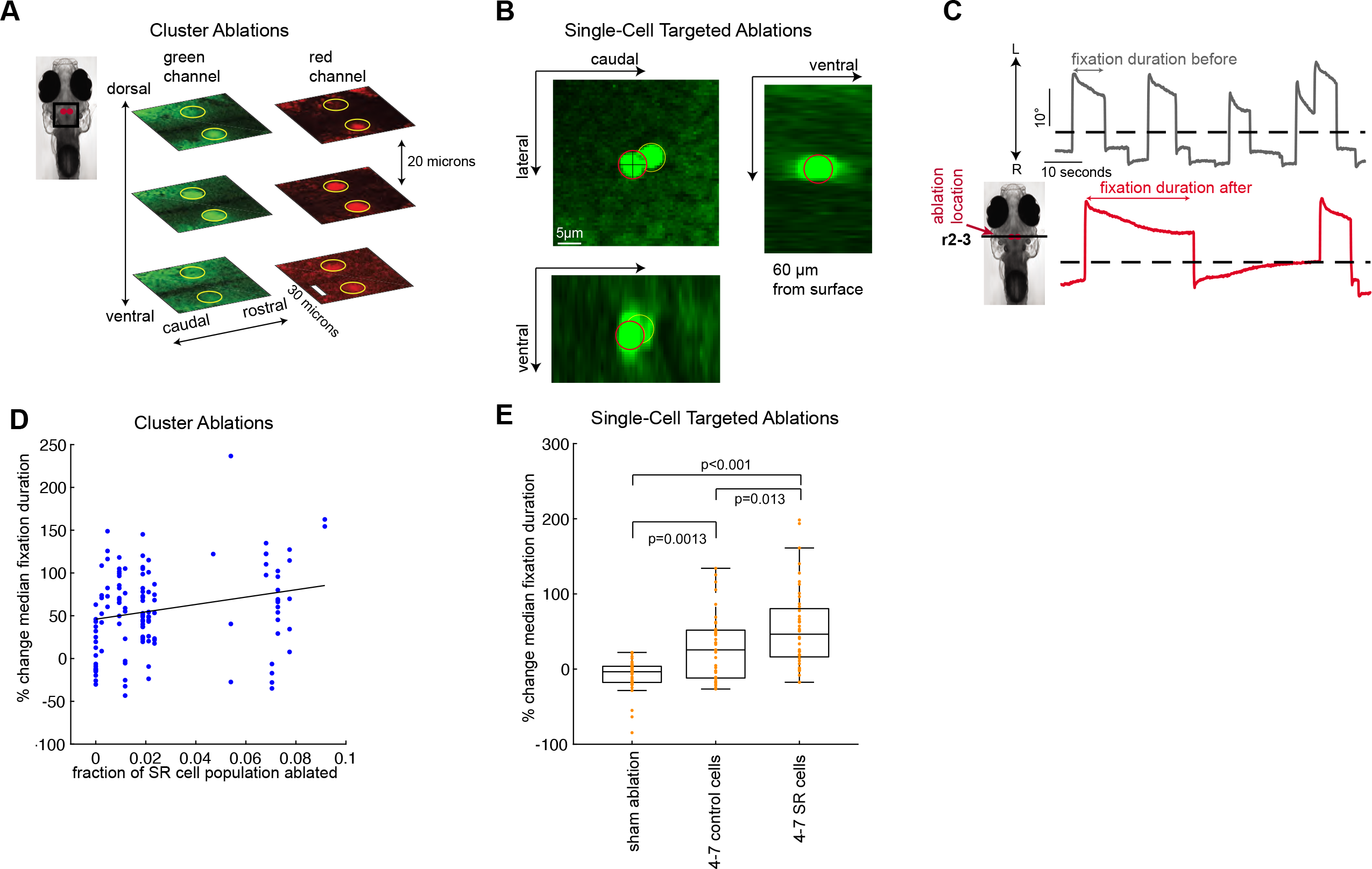
Focal laser ablations of pre-saccadic rise neurons result in increased fixation durations. (A) Time-averaged images showing horizontal sections taken from an individual fish following bilateral laser ablations of clusters of cells. Each row shows sections from dorsal (top) to ventral (bottom) locations spaced apart by 20 microns. Ablated regions appear as bright, multi-spectrum fluorescent regions approximately 30 microns in diameter (highlighted in yellow). (B) Time-averaged images taken from an individual fish following a single-neuron ablation attempt (targeted cell shown in red). Image was taken 60 microns from the surface, a typical depth where we find SR neurons. Before ablation, calcium activity and behavior were recorded and analyzed to identify SR and non-SR cells in the region. Note that both the targeted cell and an adjacent cell (outlined in yellow) were ablated. At the depths where SR cells are found, it is common for a brief high-powered laser pulse to ablate 1-3 cells. (C) Example eye position versus time recorded before (gray) and after (red) a bilateral cluster ablation was performed in the rostral hindbrain (r2-3, rhombomeres 2-3). Leftward/rightward directed saccades are made toward the direction labelled L/R. For illustrative purposes, only changes to the left eye are shown. (D) Percent change in median fixation duration as a function of estimated fraction of the SR population lost from each cluster ablation. Each blue point shows the percent change in median fixation duration following ablation in a single animal (see Methods). Pearson Correlation Coefficient between change in median fixation duration and estimated fraction of SR population lost is 0.18 (best fit line shown in black). (E) Boxplots showing percent changes in median fixation duration following single-cell targeted ablations of SR cells and control cells that did not pass our criteria for being eye-movement responsive (mean + −sem for SR targeted and control targeted ablations is 54 + −8% (n=44 from 10 animals), 28 + −7% (n=40 from 10 animals) respectively). Each orange point shows the percent change in median fixation duration following ablation in a single animal after re-sampling (see Methods). For comparison, we also show percent changes in median fixation duration from non-ablated animals using randomly selected fixations to form the sham ‘before’ and ‘after’ groups (0.27 + −2%, mean + −sem, n=41 samples from 41 animals). Central bars show the median, edges of the boxes are the 25^th^ and 75^th^ percentiles, whiskers show the range of values except outliers. Percent changes in median fixation duration in D, and E are measured as one hundred times the difference in duration (median duration after ablation minus the median duration before ablation) divided by median duration before ablation.

We found that the fixation duration generally increased following cluster ablations in the hindbrain. Since it was not clear if hindbrain ablations would affect fixation duration, we first grouped together fixations and examined changes following ablations regardless of where in the hindbrain the ablation was performed. In a separate set of control animals we performed similar size ablations in the spinal cord as a control for position-independent effects of ablation damage such as heating or vascular injury. We rejected the null hypothesis that mean fixation durations were equal before or after ablations in the hindbrain or spinal cord (one-way ANOVA, F=141.48, p < 0.01; distribution of durations shown in Supplemental Fig. 9A, mean fixation duration before and after ablations in the hindbrain equaled 12 and 17 seconds respectively (n = 8,278 fixations before and 4,101 after from 23 fish); the mean fixation duration before and after ablations in the spinal cord equaled 14 and 13 seconds (n = 911 fixations before and 1,104 fixations after from 9 fish)), and we rejected the null hypothesis that the mean fixation duration before and after hindbrain ablations were equal (difference in means and 95% CI equaled 5 [4.2, 5.7] seconds; p<0.01, two-sample t-test using a Bonferroni correction for multiple comparisons). We also tested the effect of clustered ablations on the ability of animals to maintain fixation and on saccade velocity. We found a significant decrease in fixation stability following ablations in rhombomeres 7-8, consistent with previous literature (Miri et al., 2011), but did not see a significant decrease in fixation stability after ablations in regions rostral of rhombomeres 7-8 (Supplemental Fig. 9B). We found no change in saccade velocity following hindbrain ablations (Supplemental Fig. 9C). These results provide evidence for a hindbrain role in spontaneous saccade initiation.

We found that the increase in fixation duration was correlated with the density of ablated SR neurons. We estimated the fraction of SR cells removed, for a given ablation, based on our map of their locations (Fig. 6) and plotted this fraction against change in fixation duration (Fig. 7D). The Pearson correlation coefficients between fixation duration and fraction of SR cells ablated varied between 0.01-0.35 (median cc=0.18, n=100 bootstrap samples, see Methods) and differed significantly from randomly shuffled controls (p<0.001, one-sided two-sample KS test). In a related analysis, we found that the fractional increase in fixation duration was largest for ablations in rhombomeres 2 and 3 (Supplemental Fig. 9D). In conclusion, we were more likely to see a greater increase in fixation duration when we ablated hindbrain locations that were relatively rich in SR cells compared to other regions.

Finally, we performed single-cell targeted ablations and found a significant increase in fixation duration after ablating SR neurons. In each fish, we targeted 4-7 SR cells serially, resulting in the ablation of 6-20 cells in total since 1-3 cells nearby the targeted neuron were occasionally also ablated (see Methods). To examine non-specific effects resulting from ablating a small number of cells, we targeted 4-7 neurons in a separate control group of animals that were not eye-movement responsive but were in the same region where SR cells were found. We then measured the percent increase in median fixation duration following ablation for each animal (Fig. 7E). We found SR-targeted ablations resulted in larger increases in fixation duration (45-66% [min – max], median=56%, n=100 bootstrap computations using repeated measurements from 10 fish, see Methods) than control-targeted ablations (22-39% [min – max], median=29%) and found little evidence supporting the null hypothesis that the median effect size was equal between SR and control-targeted ablations (p-values ranged from 0.0004 to 0.111, median=0.007 from 100 one-sided Wilcoxon rank-sum tests with alternative hypothesis that the median effect size in SR ablations was larger than controls, see Methods). In summary, we observed an increase in fixation duration after ablating a small number of hindbrain neurons. The magnitude of the increase was largest when we specifically targeted SR cells. Both cluster and single-cell ablation experiments suggest that SR neurons specifically play a key role in the preparation for spontaneous saccades.

## Discussion

We combined focal laser ablations and calcium imaging to comprehensively map neuronal function and activity during a self-initiated behavior. We simultaneously measured eye movements and neuronal activity throughout the hindbrain of larval zebrafish while they made spontaneous saccades in the dark. We discovered neurons in the hindbrain whose activity rises above baseline in a direction-selective manner multiple seconds before the occurrence of a saccade. We also implicated, through targeted ablations, a causal role for these cells in the decision to perform a spontaneous saccade. These data thus help elucidate the mechanism of a simple self-initiated behavior.

Our discovery of pre-saccadic (SR) ramp neurons in the hindbrain depended upon a comprehensive, single-cell resolution map of activity during spontaneous eye movements in the dark. Comprehensive spatial coverage was obtained by two-photon laser scanning microscopy, allowing us to image activity even at the deepest regions of the hindbrain where one-photon approaches suffer from poor resolution and signal-to-noise. Comprehensive analysis of the signal space was achieved by an unbiased analysis of variation that revealed three core components composing the activity of the vast majority of eye-movement related neurons. Single-cell resolution of activity was ensured by coupling two-photon microscopy with nuclear-localization of the calcium sensor, allowing us to distinguish the activity profiles of even closely packed neurons. And a focus on internally-generated dynamics was maintained by monitoring activity while animals made spontaneous saccades in the absence of any visual cues. This effort allowed us to identify both a broad spatial distribution of the various signal types and strong regional characteristics including a pronounced switch in the directional sensitivity of eye-position related signals as one crosses from rhombomere 1 to 2, a clustering of burst neurons in the ventral portions of rhombomeres 2 and 3, and a high density of SR neurons in dorsal portions of rhombomeres 2 and 3.

These mapping results complement previous work in several ways. First, in relation to a mapping effort of spontaneous activity in the dark restricted to the caudal hindbrain (Miri et al., 2011), the current map greatly extends spatial coverage but identified a somewhat sparser distributions of neurons with eye-position signals in the caudal hindbrain. This greater sparsity could be related to differences cooperativity in the calcium sensor employed. Second, in comparison to mapping work involving optokinetic behavior (Brysch, Leyden, & Arrenberg, 2019; Portugues et al., 2014), the current mapping provides information on activity associated with the preparation for and initiation of spontaneous saccadic movements. Third, in relation to a mapping effort involving phototactic behavior (Wolf et al., 2017), we observed some similarities in the distribution of position signals (e.g. the pronounced reversal in sensitivity from rhombomere 1 to rhombomere 2) but several differences like a significant clustering of neurons with pre-saccadic ramp signals in dorsal portions of rhombomeres 2 and 3 (Wolf et. al. presented preliminary findings of a region in dorsal rhombomere 7 where spatially-averaged activity rises a few seconds before exhibiting burst). This difference might have arisen because of differences in how the calcium sensor was distributed (nuclear vs cytoplasmic) and strategies for signal detection (ANOVA STA-based vs position/velocity regression-based). Fourth, in comparison to a work using locomotor behaviors to map hindbrain activity (Chen et al., 2018; Dunn, Mu, et al., 2016), the current work suggests that regions in dorsal rhombomeres 2 and 3 that have been associated with the choice of which direction the body will turn are also associated with the choice of which direction the eye will turn. Given the tendency for eye movements to precede head movements in gaze control (Freedman, 2008), it will be of interest in the future to determine what role SR neurons play in orientation behaviors more broadly. Thus, our work, together with prior mapping studies, build a solid foundation for exploring and understanding how visual and volitional signals are combined and transformed into motor commands for oculomotor behavior and potentially other orientation tasks.

Are there common underlying causes and dynamics that trigger self-initiated actions? A great number of animal movements are initiated without external cues but their causes are unknown. There is tantalizing evidence, from crayfish to humans, that a build-up in neural activity is related to self-initiated movement (Kagaya & Takahata, 2010, 2011; Maimon & Assad, 2006; Murakami et al., 2014; Park et al., 2020), but there has been no clear evidence that such neural activity is required for this type of behavior. Indeed, some experiments have suggested that buildup activity is only facilitatory, and other experiments have provided evidence that buildup activity is actually suppressive (Ames & Churchland, 2019; Lara et al., 2018; Wong et al., 2015). Our comprehensive mapping of this buildup activity enabled us to show a titration between the degree of disruption to the readiness signal and delays in the initiation of saccades. Moreover, single-cell ablations of SR neurons in particular led to significant delay in saccade initiation that was two-fold greater than any delays with ablation nearby non-SR cells. These results suggest that the spontaneous decision to move the eyes requires the build-up of activity in SR cells. Our work thus builds upon prior studies of signaling during voluntary movements by drawing a direct link between readiness signals and self-initiated movement.

The activity of SR neurons is consistent with that expected by ramp-to-threshold models for triggering behavior. In these models, a signal external to the ramp network is accumulated in value until a threshold is reached, at which point a command is given to trigger movement. While typically applied to psychophysics experiments, where the signal being accumulated represents evidence for a task-relevant decision (Churchland & Kiani, 2016; Hanes & Schall, 1996; O’Connell, Shadlen, Wong-Lin, & Kelly, 2018), the general features of such models, namely ramp rates that vary inversely with ramp times and consistent thresholds at which movement is initiated, are also applicable here. At the population level, we found that the rate of rise in SR neuron activity varied in a characteristic fashion (Figure 5D), with longer ramp times closely tied to slower ramp rates so that regardless of the fixation duration population-wide pre-saccadic activity rose to a fairly constant value at the time of saccade (Fig. 5F). Furthermore, we were able to obtain reasonable predictions as to when an upcoming saccade would occur simply by extrapolation of the ramp trend established a few seconds after SR population activity was initiated (Figure 5H). Our finding that ramp-to-threshold dynamics play a key role in the preparation for spontaneous saccades is consistent with the idea that, similar to decisionmaking tasks driven by accumulation of a signal representing external evidence (de Lafuente, Jazayeri, & Shadlen, 2015; Deverett, Kislin, Tank, & Wang, 2019), spontaneous decisions like the one studied here involve a process where some internally-generated signal that is external to the ramp network is accumulated (Murakami & Mainen, 2015; Schurger et al., 2012).

The above arguments lead to three interesting questions. First, what might the internally-generated signal(s) driving the ramp network be? Second, what is the mechanism that allows SR neurons to accumulate over time the signals they receive? Third, what might be the role of SR neurons in other saccadic behaviors such as targeted saccades and fast phase eye movements? Although these questions will require much future work to answer, several hypotheses may be attractive to investigate. Regarding the first question, one interesting possibility arises from the observation that even though neurons with fixation-related activity were widely distributed, only ablations in the caudal hindbrain led to a leaky fixation deficit characterizing loss of functionality in the velocity-to-position neural integrator (VPNI). This raises the possibility that the ongoing fixation-related activity in the rostral hindbrain could provide a kind of evidence signal that SR neurons accumulate. Regarding the second question, since the accumulation process is mathematically equivalent to temporal integration of a constant input signal, it is interesting to consider the possibility that SR neurons use a similar mechanism of recurrent excitation to promote integration as has been evidenced in the nearby VPNI for integrating saccadic inputs (Aksay, Gamkrelidze, Seung, Baker, & Tank, 2001; Joshua & Lisberger, 2015; Lee et al., 2015; Vishwanathan et al., 2017). Regarding the third question, given that SR neurons are active much earlier than midbrain and cortical cells with pre-saccadic activity (Gold & Shadlen, 2007; Hanes & Schall, 1996; Jantz, Watanabe, Everling, & Munoz, 2013) and given the widespread projections from the reticular formation to mid and forebrain areas (Moschovakis, 1997; C. A. Scudder, Moschovakis, Karabelas, & Highstein, 1996; C. Scudder, Kaneko, & Fuchs, 2002), it is possible that the SR signal in the hindbrain provides the initial kernel of activity needed to prepare movement during cued behaviors.

Answering these questions and understanding the circuit and cellular mechanisms underlying the seemingly universal ramping signal will require a combination of anatomy, perturbation experiments, electrophysiology and mathematical modeling. One key advantage of discovering the SR signal in larval zebrafish is that in this animal one has the ability to image and manipulate single cells across the entire brain (dal Maschio et al., 2017; Vanwalleghem, Ahrens, & Scott, 2018) and perform whole-circuit connectomics analysis in functionally-specified populations (Vishwanathan et al., 2017). The mechanistic insights that one can obtain on readiness in larval zebrafish will hopefully allow us to understand a broad set of decision-making processes.

## Methods

### Calcium Imaging and Eye Tracking

All experimental procedures were approved by Weill Cornell Medicine’s Institutional Animal Care and Use Committee. Transgenic larvae (7-8 days post-fertilization) expressing nuclear-localized GCaMP6f, Tg(HuC:GCaMP6f-H2B; strain cy73-431), were kindly provided by Misha Ahrens’ lab. Fish were embedded in 1.5% low-temperature agarose and subsequently imaged using a custom-built two-photon laser scanning microscope (Daie, Goldman, & Aksay, 2015). We performed two-photon imaging using excitation light (930 nanometers) from a tunable laser (Spectra-Physics Mai Tai) sent through a 40x (0.8 NA) water-immersion objective lens (Olympus LUMPLFL40XW/IR2) to the hindbrain. The laser power was controlled using an electro-optical modulator (Conoptics 350-50UV) and amplifier (Conoptics 302RM). Laser power used for imaging ranged from 15-25 mW at the sample. Neurons within square horizontal planes (185 microns in length; 5-20 planes per fish spaced apart by 5 microns) were imaged simultaneously at 0.98 Hz (512 lines at 2 milliseconds per line). Recordings lasted 4-5 minutes per plane. Images were saved as uncompressed grayscale TIFF stacks. Horizontal eye movements were tracked in real time using a sub-stage CMOS camera (Allied Vision Technologies, Guppy FireWire camera) illuminated with infra-red light (850nm, Thorlabs 851L). Custom Matlab software extracted horizontal eye position (e in Figure 1A) from camera images in real-time at a variable sampling rate of approximately 13 Hz based on an algorithm described in (Beck, Gilland, Baker, & Tank, 2004; Miri et al., 2011).

### Registration of Individual Planes to the Z-Brain Atlas

Each imaging plane was first registered to a corresponding plane in a reference “bridge” brain constructed from a single Tg(HuC:H2B-GCaMP6f) fish (8 dpf) using two-photon microscopy. Single planes in the bridge brain was constructed by stitching together six overlapping 512×512 (185 micron squared area) images through an optimized translation procedure using the *FIJI* Grid/Collection Stitching Plugin (Preibisch, Saalfeld, & Tomancak, 2009; Schindelin et al., 2012). Each image used in the stitching procedure was formed by averaging calcium activity over a 20 second interval. The six images used were sufficient to completely tile any horizontal plane in the hindbrain. Bridge brain images were taken from planes (spaced apart by 3 microns in the dorsal-ventral axis) that extended from the most dorsal portion of the hindbrain to the most ventral portion where expression was no longer visible. For each animal, we computed a 4×4 affine transformation matrix that linearly transformed 3-dimensional (3D) points from that animal to 3D points in the bridge brain. We used the *BigWarp* tool (Bogovic, Hanslovsky, Wong, & Saalfeld, 2016) in *FIJI* to select several 3D points, *x*, based on features such as fiber bundles, the midline, and ventricles in each imaged animal and to match these points with corresponding 3D points, *y*, in the bridge brain. The affine matrix was calculated in Matlab using the ‘\’ operator to find a matrix, *T*, that solved the equation [*x*, 1]*T* = [*y*, 1]. The fourth column of *T* corresponding to translations was then set to equal 1 in the fourth row and zero at all other rows. We repeated this procedure to find corresponding points and an associated transformation matrix between the bridge brain and the Elav3-H2B brain available on the Z-Brain website. We qualitatively inspected each registration using the output from the *imwarp* function in Matlab. If necessary, we selected additional 3D points (at least 6) until results were visually similar to the example shown in Supplemental Figure 1A.

### Automated Identification of Regions of Interest

Prior to analysis, each calcium movie was pre-processed to correct for motion artifacts (see Supplemental Procedures for details).

#### Finding the Locations of Active Cells Based on Fluorescence Dynamics

We implemented the alternating matrix factorization algorithm described in detail in (Pnevmatikakis et al., 2016) on motion-corrected fluorescence movies using the freely available CalmAn-MATLAB matlab toolbox provided by the Flatiron Institute (https://github.com/flatironinstitute/CaImAn-MATLAB). Briefly, the algorithm models a calcium fluorescence movie as the product of two non-negative matrices, one containing spatial locations and the other containing calcium time-series for each active cell, plus a background and noise component. To determine the non-negative matrices that best-fit the data, the algorithm performs multiple iterations of constrained least-square optimizations for one of the two non-negative matrices while holding the other non-negative matrix fixed. The matrix being fit to the data and the matrix that is held fixed alternate with each iteration. We implemented a procedure based on the ‘*demo_script.m*’ file provided with the code. After initializing the spatial and temporal components with a greedy procedure, we ran 1 iteration of spatial and temporal updates, followed by a post-processing step where similar components were merged and unrealistically large components were removed, followed by a final spatial and temporal update. We investigated different values of the number of neurons parameter, *K*, and chose a value (250) that produced reasonable looking spatial footprint estimates based on visual inspection of a handful of sample fluorescence movies. Fluorescence traces used in subsequent analyses were found by spatially averaging across pixels within the final spatial footprints found by the above procedure (see Supplemental Procedures for more details on the CalmAn-MATLAB routines used). We registered each cell to a reference brain (see Methods *Registration of Individual Planes to the Z-Brain Atlas*) and excluded cells that were registered to the midbrain.

In the results (*Mapping Activity Using Two-Photon Microscopy*) we give estimates of the total number of cells in our data set and in Supplemental Figure 1D, 1E we make reference to non-active cells. For these sections only, we also relied on a separate procedure to infer cell locations (see Supplemental Procedures-*Measuring the Total Number of Cell Nuclei*) since the alternating matrix factorization algorithm cannot find non-active cells (non-active cells are included as part of a single background term).

### Automated Determination of Saccade Times

We determined the times of saccade occurrence by calculating the crossing times of eye velocity past a threshold. To calculate eye velocity, we first filtered out fluctuations in eye position using a median filter (*medfilt1* in Matlab 9.5.0.9444). The exact value of the filter order depended on the eye position sampling rate but was chosen to correspond to 500 milliseconds. We then approximated eye velocity as the difference in filtered eye position at consecutive time points divided by the time difference between these points. The threshold was set to three standard deviations above the mean absolute velocity or 10 degrees per second, whichever was larger. A single saccadic event typically consisted of several consecutive points whose velocity was above the threshold. We took the initial point as the time when the saccade occurs. During head/body movements the eye position traces become corrupted. One signature we use to determine when head/body movements occur is the time between threshold-crossing events; separate experiments with video recordings of the entire body suggested that unusually short intervals between events typically indicate that the events occur during sudden head/body movements. For this reason, we did not consider threshold-crossing events that were spaced apart in time 1.4 seconds or less to be saccades (see Supplemental Experimental Procedures for additional calcium imaging based criteria for detecting head/body movements).

### Saccade-Triggered Average (STA) Calculation

We averaged saccade-triggered signals across fixations after linearly interpolating them to a grid of time points equally spaced apart by 1/3 seconds. dF/F was computed as (F-F0) / F0 where F is the average fluorescence within a cell’s ROI and F0 is the mean fluorescence across the entire recording period. dF/F before and after each saccade was extracted, interpolated (using *interp1* in Matlab 9.2.05 with the *linear* method), grouped across saccades and then averaged according to the direction of saccade (left or right). We excluded any cells that were recorded during an experiment that contained less than five saccade-triggered responses total (either saccades to the left or right). The amount of time around saccade we could examine was directly related to the number of samples available to select cells related to eye movements; if a group of cells were recorded when the fish made few long-lasting fixations then these cells needed to be excluded from further analysis since their STAs would be undefined at long times away from saccade. We chose a window that extended five seconds before and after saccade because at this value we retained a reasonable amount of data while still finding patterns with time across the STAs (Figure 3C). 95% confidence intervals about the average (Figure 2B) were found by resampling the saccade-triggered responses with replacement and calculating the STA for each resample (number of resamples = 100). We measured the lower and upper bounds of the confidence intervals as the 2.5% and 97.5% quantiles across the bootstrapped samples.

### Selection of Cells Related to Eye Movements

We ran a One-Way ANOVA (Matlab 9.2.05 function *anova1*, ‘Statistics and Machine Learning Toolbox’) on saccade-triggered responses to test the null hypothesis that STA activity was equal at all timepoints versus the alternative that at least one timepoint had average activity that differed from the others. We considered a neuron as being eye movement-related if we rejected the null hypothesis for either one of that neuron’s two STAs (the STA triggered around saccades to the left or right). We used the Holm-Bonferroni method (Holm, 1979) to correct for multiple comparisons. This procedure varied the significance level for each comparison using the formula p/(N – j +1), where j was the index of the comparison after sorting p-values from low to high, p was the desired family-wise error rate, and N was the total number of comparisons. We set p to 0.01 and set the number of comparisons to 63,644*2=127,288, which was the total number of hindbrain ROIs found by the NMF algorithm times two to account for both saccade directions.

### Principal Component Analysis

We ran a principal component analysis (PCA) to search for lower-dimensional representations of the STAs across the population. We combined STAs from all cells (across all planes and fish recorded) and from both directions (around saccades to the left and right) resulting in a matrix, *f_i_*(*t_j_*), analyzed by PCA that had 13,800 samples (*i* =1,…,13,800; 6,900 cells times two directions) and 31 dimensions (time around the saccade is evaluated at 31 discrete time bins, *t_j_* for *j* = 1,…,31, covering 10 seconds at 1/3 second binsizes). PCA (computed using Matlab 9.5.0.9444 ‘Statistics and Machine Learning Toolbox’ function *pca*) resulted in a set (31 in total) of 31-dimensional orthonormal basis vectors, 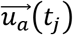, for *a* = 1,…,31, which we refer to as components (Figure 3A) and a matrix of coefficients, *c*, (Figure 3Bi) that, after subtracting the population-mean from each row, characterize the matrix of STAs by the equation

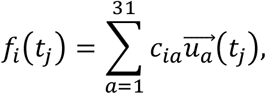

for *i* = 1,…, 13,800, *j* = 1,…,31. Note that the coefficients determine how much each component, 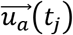, contributes to an individual STA. Since the elements of the components carry the time-dependence of the STA response and since the components are orthogonal, the magnitude of the coefficient vectors formed by the rows of *c* determine the, time-independent, magnitude of the STA responses. After finding that most of the variance across the population could be explained by the first three components (Figure 3A), we focused on the coefficients corresponding to these components. We examined normalized coefficients, 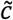, computed by dividing the coefficients by root sum of squared values across the first three components

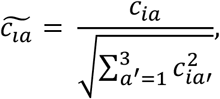

where the coefficients have been sorted so that *c*_*i*1_ corresponds to the component explaining the most variance, *c*_*i*2_ corresponds to the component explaining the second most variance, and so on. The principal advantage of this normalization is that we can examine the distribution of coefficients in two-dimensions by only sacrificing information about the magnitude of STA responses since the vectors formed by the rows of 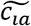 have unit norm. If we only examined the non-normalized coefficients of the first two components we would also sacrifice information about the temporal variations of STA responses. Throughout the text we work with the normalized coefficients converted into spherical coordinates *ϕ* and *ϑ* (see Figure 3Bii)

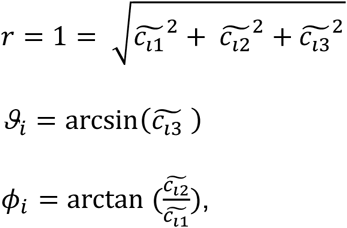

for *i* = 1,…, 13,800. For visualization purposes, the values of *ϕ_i_* were then shifted to lie between −180 and 180 degrees, and the values of *ϑ_i_* were defined to be 0 degrees at the equator of the globe in Figure 3Bii, iii (90 degrees minus the azimuthal angle in spherical coordinates) and then shifted to lie between −90 and 90 degrees as shown in Figure 3Biii. The two-dimensional probability density functions over *ϕ* and *ϑ* shown in Figure 3Bii, iii were measured using a normal Gaussian kernel smoothing function (Matlab 9.2.05 ‘Statistics and Machine Learning Toolbox’ function *ksdensity* with bandwidth parameter equal to 10 degrees for phi and 3 degrees for theta). Note that the population averages shown in Figures 3Biv, 3C and Supplemental Figures 3A, 4 are constructed from responses that are ordered by the normalized coefficients *ϕ* or *ϕ* and *ϑ* but are not constructed by linearly summing the first three components weighted by the normalized coefficients. Therefore, these population average responses are not unit normalized and can reflect variations not captured by the first three components.

### Anatomical Projections of Normalized Coefficients Derived From PCA

We created maps of *ϕ* (see Figure 3Bi, 3Bii and methods section *Principal Component Analysis* for a definition of *ϕ*) for saccades to the left/right for each fish and imaging plane. For each map, we set the value at an individual cell’s location to the value of *ϕ* around saccades to the left (Figure 3D)/right (Supplementary Figure 3B) measured for that cell. Individual maps were then registered to a reference brain using the procedure described in the methods section *Registration of Individual Planes to the Z-Brain Atlas*. The values of *ϕ* from individual cells registered to overlapping three-dimensional locations were averaged together and regions of space without any cells were set to NaN. The resulting three-dimensional map of *ϕ* was used to create the average projections shown in Figure 3D/Supplementary Figure 3B. To avoid viewing regions showing a cell that only appeared in one fish, we removed regions where the ratio of cells to fish in the average projection was less than 0.5.

### Distinguishing SR Cells from Non-SR cells

We calculated the correlation coefficient between the time until an upcoming saccade and dF/F before each saccade using Spearman’s rho (Matlab 9.2.05 ‘Statistics and Machine Learning Toolbox’ function *corr* with option ‘type’ set to ‘Spearman’). The Spearman correlation can be used to measure monotonic (not only linear) relationships between two variables X and Y. It is calculated as the standard Pearson correlation coefficient applied to the ranks of X and Y. We did not interpolate fluorescence activity before computing the correlation coefficient. We measured a correlation coefficient for each eye movement-related cell. We computed a p-value for each correlation coefficient by testing the hypothesis that rho = 0 against the alternative that the correlation was greater than 0 (‘tail’ option set to ‘right’). A neuron was considered to have significant pre-saccadic activity if we rejected the null hypothesis before either leftward or rightward directed saccades at a significance level of 0.01. We used the Holm-Bonferroni method to correct for multiple comparisons.

### Measuring Time and Rate of Pre-Saccadic Rise

#### Measuring Time of Pre-Saccadic Rise

We estimated the timing of pre-saccadic activity in SR cells by first constructing non-negative, deconvolved estimates of their activity (see Supplemental Procedures – Constructing NonNegative Estimates of SR Activity). The non-negativity constraint resulted in epochs of time where the deconvolved estimate was equal to or nearly zero which we interpreted as epochs where the cell was inactive. Based on visual inspection of deconvolved SR traces, we could reasonably distinguish times when activity crossed from inactive to active, and vice-versa, by determining when deconvolved activity crossed a threshold near zero (0.1). Therefore, we measured the time of pre-saccadic rise as the time corresponding to one sample before deconvolved SR activity increased above 0.1. Brief (1-2 sample) dips below zero in inferred spiking activity occasionally occurred and were not counted as threshold crossing events.

#### Measuring Rate of Pre-Saccadic Rise

We estimated the rate of pre-saccadic rise in SR cells by finding the slope of the best-fit line of pre-saccadic dF/F activity with time. To construct the best-fit line we used time from pre-saccadic rise to the time of upcoming saccade as a regressor to a linear regression that fit dF/F values. The linear approximation was reasonable for 89% of the fixations (correlation between regression fit and data was greater than 0.4). We excluded fixations where we were unable to measure the slope with linear regression.

### Saccade Time and Occurrence Predictions Based on Pre-Saccadic Activity

We predicted whether a saccade will occur and when a saccade will occur based on interpolated SR activity before saccadic events. Throughout this section, the phrase *interpolated SR activity* refers to linear interpolation of dF/F activity to a grid of equally spaced time points (using 1/3 second bins) starting from the previous saccade to the upcoming saccade.

#### Predicting Whether a Saccade Will Occur Using Choice Probability

The choice probability (CP) is a commonly used metric in neurophysiology and psychophysics experiments that quantifies how well an ideal observer can predict animal behavior (Crapse & Basso, 2015). In a typical neurophysiology application, the CP measures relationships between neuronal discharges and binary behavioral choices. We adapted this metric to our setting by using the spontaneous decision to saccade to the left or the right as our binary behavioral variable and population average dF/F at a given time before saccade as our neural read-out. At discrete timepoints before upcoming saccades, we made two histograms of interpolated SR activity. One distribution, the “noise” distribution, was comprised of values of dF/F before saccades to the right (left) from cells whose dF/F was significantly correlated with upcoming saccades to the left (right). The other distribution, the “signal” distribution, was comprised of values of dF/F before saccades to the left (right) from cells whose dF/F was significantly correlated with upcoming saccades to the left (right). The decision to saccade was predicted using a threshold on population activity. We plotted the fraction of dF/F values from the “signal” distribution that are above threshold (the true positive rate) versus the fraction of dF/F values from the “noise” distribution that are above threshold (the false positive rate) across multiple threshold values. The CP was calculated as the area under the resulting curve, which would equal 0.5 if dF/F and upcoming saccade choice were not related.

#### Predicting Saccade Times Using a Ramp-to-Threshold Model

We predicted the saccade times by calculating a running prediction of population activity slope and then using this estimate to determine when population activity will cross a threshold, *θ*. We first measured *θ* by averaging dF/F reached at the time of saccade, relative to pre-rise baseline, across all cells, fish, and fixations. For each fixation duration, we then calculated population average activity before upcoming saccade by combining (across all cells and fish) interpolated SR activity (we used eighteen non-overlapping values of fixation duration from 3.5 + −0.5 seconds to 20.5+ −0.5 seconds). We modeled the population average dynamics, *y*′(*t*), at values of time *t* after population activity begins to rise, with the following linear ramp equation

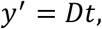

where *D* is the population activity slope. After a time, *t_r_*, a saccade will occur and *y*′ should equal *θ* if the ramp-to-threshold model is accurate, e.g.

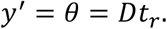

We constructed a running estimate of the slope by first measuring when the actual population activity, *y*(*t*), began to rise. We measured this as the time when the derivative in *y*(*t*) crossed a threshold of 0.04 (dF/F)/s. Note that under our convention we set this event to occur at time 0. The derivative was approximated as the difference between population activity at each time point divided by the interpolated time bin interval, Δ*t*, of 1/3 seconds. We created a running estimate of the population activity slope, which we denote as 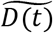, using the median value of the derivative from time 0 until time, *t*,

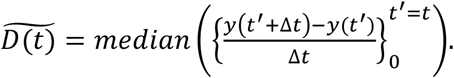

We substituted our running estimate of the slope into this equation to yield a running estimate of *t_r_*

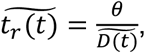

where we have used the ~ symbol to distinguish our running estimates from the actual values. Using our estimate of 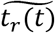 we could predict the amount of time until upcoming saccade for any given value of *t*, as 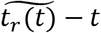. Figure 5H shows predictions of the time until upcoming saccade made by varying the value of *t* from one time point after population activity begins to rise to one time point before saccade for each fixation duration. In the text and in Supplemental Figure 7, we also present results showing how well the population average activity is approximated by a ramp-to-threshold model. In these cases, we did not create a running estimate of the slope but instead only created one slope estimate by setting (*t* = *t_r_*) in the above equation for determining 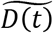. We used all cells to estimate 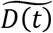 when we forecasted saccade times for Figures 5H, 5I, however using all cells would have led to overfitting in Supplemental Figure 7 since we base population average estimates on all time points. To prevent overfitting in this case, we only used a random subset of all cells (60%) to create *y*(*t*), 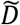, and the resulting ramp-to-threshold model, *y*′(*t*). We then created a new estimate of population average activity using the remaining 40% of cells to test against *y*′. To determine model accuracy, we repeated this procedure on 10,000 randomly selected subsets from randomly chosen fixation durations.

#### Saccade Time Predictions by an Ideal Observer

As a control, we determined the fraction of saccades that could be accurately predicted given knowledge of the elapsed time since last saccade and the distribution of fixation durations. Given this information, an ideal observer could predict upcoming saccade times by guessing a time that minimizes some cost-function that measures error between the actual saccade time and the guessed time. We tried three cost functions (mean-squared error, mean-absolute deviation, and all-or-none error) and report results in the text from the one that performed the best (all-or-none error).

### Laser Ablations

Ablations were performed with the same microscope used for imaging.

#### Cluster Laser Ablations

We targeted regions along the dorsal-ventral axis that were 30 microns dorsal of the medial longitudinal fasciculus since this is where we found the majority of eye movement-related cells. Cluster ablations were created by implementing the procedure described in (Miri et al., 2011), we focused the laser to an area smaller than a single cell and then increased the average laser power to values between 130-150 mW for 1-5 seconds. We repeated this procedure until we saw a lesion, which we determined by looking for a multi-spectrum spot that was much brighter than the fluorescence of surrounding tissue. Lesion sizes with this procedure were generally around 5 microns in diameter. To increase the size of the lesion we lowered the average laser power to values of 30-50 mW and scanned the ablated region at these lower powers which caused the lesion size to grow. We stopped scanning the ablated region once it grew to approximately 30 microns in diameter. We waited between 30-120 minutes after ablation before recording post-lesion eye-movements.

To estimate the fraction of SR cells lesioned in a given fish during cluster ablations (Figure 7D and text), we calculated the number, nc, of SR cells from the 3D map of SR locations (whose projections are shown in Figure 6) that fell within a cylinder (30 micron radius, 60 micron side length along dorsal-ventral axis) centered at the registered lesion location. To determine the cylinder center, we manually inspected time-averaged images of each the ablated region to find planes containing a bright, multi-spectrum fluorescence characterizing ablation damage. We found the plane with the maximal damage, selected similar points between this plane and the bridge brain (as demonstrated in Supplemental Figure 1A) and then created a 2D affine matrix to transform the center of the registered lesion to the bridge brain. The fraction ablated was equal to nc divided by the total number of SR cells used to construct the map.

#### Single-Cell Laser Ablations

In a separate set of experiments, we targeted neurons for ablation individually after checking for pre-saccadic ramping activity. Before these ablations, we first processed calcium movies during the experiment to identify the locations of SR neurons and cells not related to eye movements, which took between 10 minutes to two hours. After processing, we zoomed in to the region with the cell of interest, took a time-averaged image, and manually corrected the location for any changes in cell-center location that might have arisen during processing. We did not attempt an ablation if the experimenter could no longer manually identify the cell of interest after processing. Immediately after this step we ablated individual neurons by focusing a high-powered, pulsed femtosecond laser (810nm, 400-500mW after the objective) on the center of SR cells for a brief (2-5 ms) period of time (Koyama et al., 2016; Liu & Fetcho, 1999; Vladimirov et al., 2018). This procedure resulted in the loss of 1-3 cells per ablation attempt (Figure 7B) at the depths where SR cells were targeted (40-70 um below the surfaced). In some cases, ablations did not occur even after 3-5 attempts, most likely due to laser power absorption from pigmentation. We did not try to ablate a cell after 5 attempts. In one experiment, we repeated three cycles of finding cells of interest in a single plane, ablating these cells, then searching for more cells of interest in subsequent planes. In all other experiments we imaged, identified cells of interest, and then ablated these cells.

#### Analysis of Behavior Following Laser Ablations

For each fish, we measured fractional changes in median fixation duration after ablation. Since we were concerned with a spontaneous behavior, we could not control the number of fixations that were recorded during the timeframe of each experiment. As a result, our measurements would have had different accuracies per animal (Supplemental Table 1) if we calculated median changes without any procedure to account for the different numbers of fixations observed per fish. To control for this difference in accuracy while using all of our data, we made repeated measurements of the fractional change per fish with each repeated measurement having the same number of fixations before and after ablation. For each repeated measurement, we randomly sampled without replacement *N_min_* fixations before and after ablation where the exact value of *N_min_* differed between single-cell and cluster ablation experiments and was based on animals with the fewest number of fixations available. Specifically, if we denote the number of fixations in condition *i* (where *i*=1 denotes before ablation and *i*=2 denotes after ablation) from animal *j*, as *n_ij_*, then 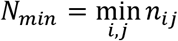 (*N_min_* equals 33 fixations for single-cell ablation experiments and 32 fixations for cluster ablation experiments). The number of times we repeated each measurement of fractional change in median fixation duration varied per fish and was determined by how many more fixations each animal made compared to *N_min_*. Specifically, the number of times we repeated each measurement for animal *j* equaled 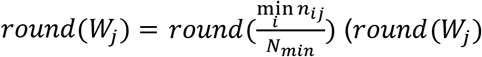 varies between 1-11, see Supplemental Tables 2, 3 for exact values per fish). The fractional change in median fixation duration was computed as the difference in median fixation duration (after minus before ablation) divided by the median fixation duration before ablation. Using *l* to denote the index of the repeated measurement in animal *j* and *t_ijml_* to denote *mth* randomly sampled fixation duration (integer *m* varies from 1 to *N_min_*) in this measurement in condition *i* (before or after ablation), the fractional change in fixation duration was computed as:

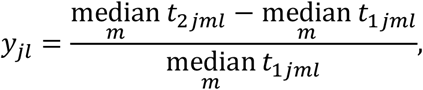

for *l* = 1,2,…, *round*(*W_j_*) and *j* = 1,2,…, total number of animals (see Supplemental Tables 2, 3 for total number of animals per group). Since the set of measurements *y_jl_* and our subsequent results depend on the random samples chosen, we repeated our sampling procedure 100 times to obtain 100 different sets of measurements *y_jl_*. The values plotted in Figure 7D, 7E, Supplemental Figure 9D show the output from one of the 100 runs performed. In the results section regarding cluster ablations (see *Focal Laser Ablations Identify SR Cells as Indispensable for Setting Spontaneous Fixation Durations*), we presented the minimum, maximum, and median from 100 calculations of correlation between the points *y_jl_* and the associated estimated fraction of cells removed during a cluster ablation (the fraction of cells removed was calculated once for animal *j* and did not vary with measurement *l*). Figure 7D shows one run of *y_jl_* versus fraction of cells ablated that results in a typical correlation coefficient; Supplemental Figure 9D shows results from the same run used to construct Figure 7D. In the results section regarding single-cell targeted ablations, we calculated the mean across *j* and *l* of *y_jl_* for each run and presented the minimum, maximum and median result (Figure 7E shows one run of *y_jl_* that results in a typical change between SR and control targeted groups; the sham ablation results presented in Figure 7E were computed without making repeated measurements using *N_min_*=33 fixations per animal to compute fractional change in median fixation duration). For each set of *y_jl_* for control and SR-targeted animals, we ran a Wilcoxon Rank-Sum test (100 tests in total) of the Null hypothesis that the medians are equal between the distribution of *y_jl_* for SR-targeted and the distribution of *y_jl_* from control animals against the alternative that the median is greater SR-targeted animals. The minimum, maximum, and median p-values across the 100 runs were presented. Matlab function *ranksum* with the appropriate value of ‘*tail*’ was used to perform the one-sided Wilcoxon tests.

For more details regarding imaging analyses and statistics see Supplemental Experimental Procedures.

## Data Availability Statement

The data used in this study are available from the corresponding author upon reasonable request.

## Code Availability Statement

The code used for data analysis in this study is available from the corresponding author upon reasonable request.

## Acknowledgements

We thank Chao Young and Misha Ahrens for generating the Tg(HuC:GCaMP6f-H2B) line and sharing ahead of publication. We thank the RARC team at Weill Cornell Medicine for animal care and husbandry. We thank Misha Ahrens, Ari Arrenberg, Robert Baker, Tom Chartrand, Chanwoo Chun, Mark Goldman, Charlotte Grove, Minoru Koyama, Gaby Maimon, Sandro Romani, and Karel Svoboda for helpful discussion and feedback on the manuscript. Funding for this work was provided by NIH grant K99 EY027017, R01 EY021581 and the Simons Foundation Global Brain Initiative.

## Author Information

A.D.R, and E.R.F.A. conceived the project; A.D.R carried out the experiments and analyzed the data; A.D.R and E.R.F.A wrote the paper.

## Ethics declarations

The authors declare no competing interests.

## Supplemental Experimental Procedures

### Image Pre-Processing to Correct for Small Movements

We were able to correct for small drifts that occurred during imaging using a motion-correction procedure based on cross-correlation. We first calculated the median fluorescence intensity across time for each pixel in the movie and used the resulting image as a reference. Any frame in the movie that deviated from the reference image was considered to have a movement artifact which required correction. To register each movie frame to the reference, we used the Matlab function *dftregistration.m* which implemented the two-step discrete Fourier transform (DFT) algorithm described in (Guizar-Sicairos et al., 2008). For computational efficiency the *dftregistration* algorithm worked in Fourier space to calculate cross-correlations. We used Matlab’s built-in fast Fourier transform software (*fft2*) to compute each frame’s two-dimensional DFT and to compute the two-dimensional DFT of the reference frame. Briefly, the *dftregistration* algorithm estimated the peak in the two-dimensional cross-correlation between the reference image and movie frame being registered. Each movie frame was then translated by an amount determined from the peak location.

### Detecting Samples Corrupted by Animal Movement

The motion-correction algorithm described above returned a scalar metric for each movie frame that indicated how well the frame matched the reference after correction. If this value was too low, the frame was considered too aberrant to be useful and the fluorescence of all pixels in this frame were replaced by NaNs. Specifically, for each frame, the *dftregistration* algorithm returned an error value related to the square root of one minus peak, normalized cross-correlation between a given frame and the reference. For each imaging plane, we computed the median error across all frames and the median absolute deviation (MAD) of the error across all frames. If a given frame’s error value was greater than 5 times the MAD plus the median that frame’s pixels were replaced by NaNs.

### Measuring the Total Number of Cell Nuclei

We analyzed time-averaged images to estimate the total number of neurons in our dataset. For each motion-corrected fluorescence movie, we calculated the median intensity across time for each pixel and analyzed the resulting time-averaged image to find individual cell nuclei locations. We performed a morphological opening on the time-averaged image (Matlab function *imopen*) with a disk-shaped structuring element that had a radius equal to 4 pixels (1.4 microns). The opening operation with this structuring element tended to make it easier to segregate the disk-shaped nuclei in the image. We then found local intensity maxima of the opened image by looking for connected pixels with equal intensity that were greater than the intensity of external boundary pixels (Matlab function *imregionalmax*). We measured the locations of individual cell nuclei measuring the regions of connected pixels that corresponded to local intensity maxima. To control for false positives, we excluded any selected ROIs that had an area greater than typical cell areas that we found by manual measurements (19 microns squared which translated to 144 pixels squared).

### CalmAn Routines for Locating Cell Nuclei

We first found initial estimates for the spatial, temporal and background components using the *initialize_components* function. This function ran several steps: 1) it spatially filtered the fluorescence movies (Gaussian kernel with standard deviation set to 5, which corresponds to 1.8 microns). 2) it greedily selected locations where the spatial estimates explained the largest amount of spatio-temporal variance. 3) it used rank-1, non-negative matrix factorization to produce spatial, temporal, and background estimates. 4) it refined these estimates using a hierarchical, alternating, non-negative matrix factorization method. 5) it ran a rank 1 non-negative matrix factorization on the spatio-temporal residual to initialize the background spatial and temporal components. We updated the initial estimates of the spatial footprints and the background component using the constrained non-negative Lasso algorithm implemented in the *update_spatial_components* function. We used the ‘dilate’ option which computed the search location by dilating the support set of each component from the previous iteration using a 4-pixel radius (1.4 microns) disk-shaped structuring element. The new components are then post-processed by the following operations: (i) 2-d median filtering with a default size of 3×3 pixels (ii) morphological closing with a length 3 pixel, square shaped structuring element, and (iii) energy thresholding with threshold set to 0.99. We then updated the estimates of the temporal components using the *update_temporal_components* function with an auto-regressive parameter, p, equal to zero. This function updated components using a block-coordinate descent algorithm (we used 2 iterations) which, with p equal to zero, ran a thresholding operation (at a threshold of 0) on the activity of each component after removing the effect of all the other components. After one spatial and temporal update, we removed spatial or temporal components that were poorly correlated with the raw data (space and time r-values returned by *classify_comp_corr* function were less than 0.05) or whose spatial footprints were larger than a threshold (16 pixels which equaled 5.8 microns). We then merged spatially overlapping components with highly correlated temporal activity (cc>0.95) using the *merge_components* function. Using the new component estimates we then ran one more iteration of the spatial and temporal component updates described above. Finally, since we were only interested in the spatial locations of active neurons, we set non-zero values for each spatial component to one and estimated each component’s neuronal activity as the trace that resulted from spatially-averaging fluorescence at non-zero locations.

### Constructing Non-Negative Estimates of SR Activity

We created non-negative, deconvolved estimates of SR activity using the *update_temporal_components* function provided by the CalmAn-MATLAB matlab toolbox. The function *update_temporal_components* provided estimates of denoised fluorescence activity and non-negative spike estimates given spatial components, a background signal, and parameters for the calcium impulse response function. For each SR cell, we ran *update_temporal_components* with the auto-regressive parameter set to one (corresponding to a single exponential calcium impulse response function). We fixed the auto-regressive time constant to 1.8 seconds (in the equivalent continuous-time model) which roughly corresponded to the calcium buffering time for nuclear-localized GCaMP6f (Kawashima et al., 2016). We used the spatial components of SR cells found by our original run of the alternating matrix factorizations procedure as described in section CalmAn Routines for Locating Cell Nuclei. For the spatial background component we used the time-averaged image of the plane containing the SR cells being analyzed with holes (zero values) at the SR cell locations (determined by their spatial footprint). We used the average across all pixels as the temporal background component. As described in the documentation, *update_temporal_components* ran a block-coordinate descent algorithm (we used 2 iterations), which deconvolved the activity of each SR cell after removing the effect of all the other SR cells. We used the default ‘*constrained_foopsi*’ method along with the CVX toolbox for deconvolution, which solved a noise-constrained optimization problem to produce estimates of denoised fluorescence activity and non-negative spike estimates. The noise level for each neuron was estimated by averaging the power spectral density of component fluorescence activity over “high” frequencies (one half the Nyquist frequency to the Nyquist frequency which equaled 0.24-0.49 Hz for our data).

**Supplemental Table 1:**
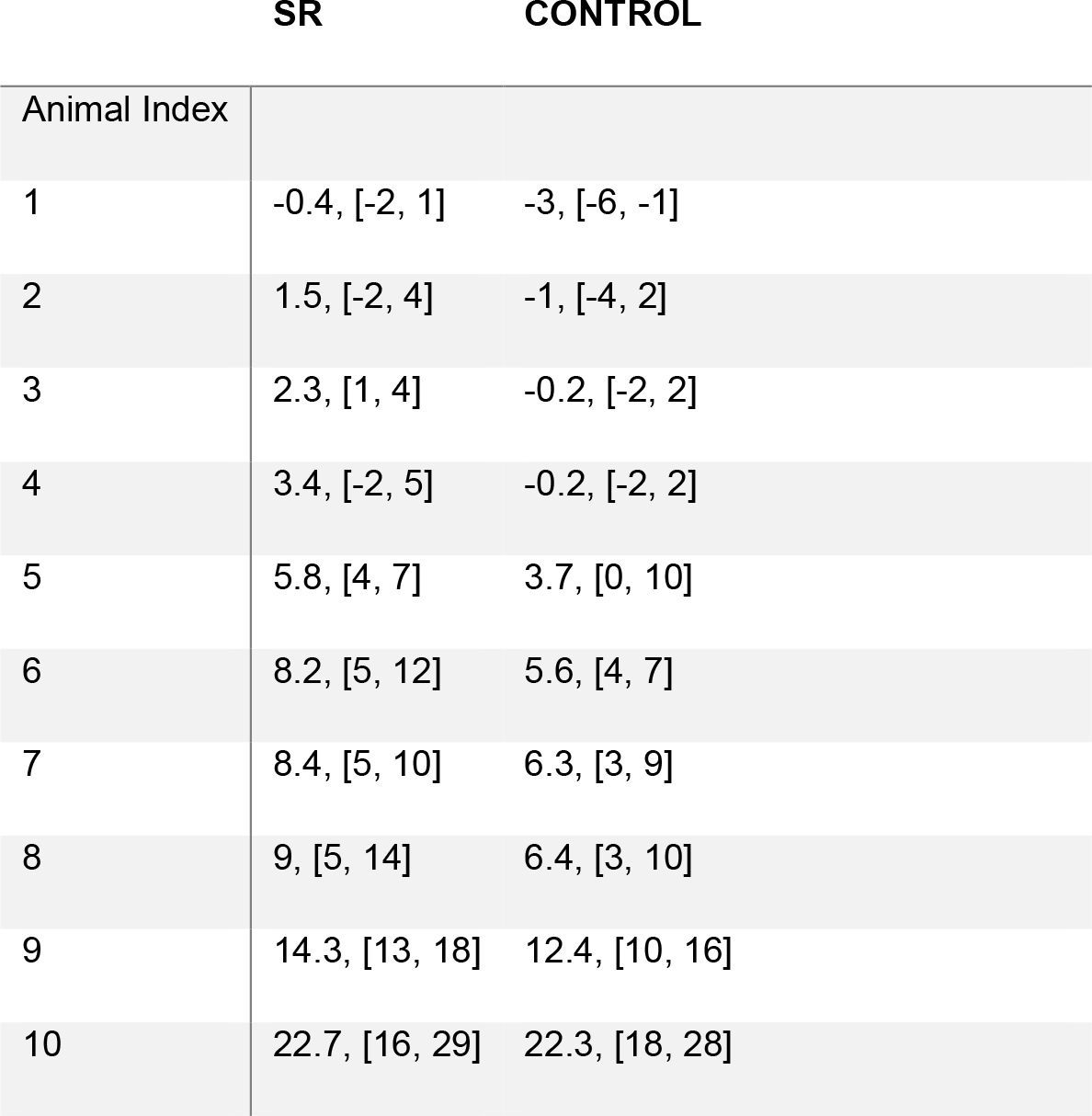
Increase in median fixation duration along with 95% confidence intervals (in brackets) following single-cell ablations without controlling for differences in number of fixations per animal. Notice how the variability changes per fish. Values are reported in seconds. Confidence intervals were found by bootstrapping with 10,000 samples. Note that animals with SR-targeted ablations are different than animals with control ablations.

|  | SR | CONTROL |
| --- | --- | --- |
| Animal Index |  |  |
| 1 | -0.4, [-2, 1] | -3, [-6, -1] |
| 2 | 1.5, [-2, 4] | -1, [-4, 2] |
| 3 | 2.3, [1, 4] | -0.2, [-2, 2] |
| 4 | 3.4, [-2, 5] | -0.2, [-2, 2] |
| 5 | 5.8, [4, 7] | 3.7, [0, 10] |
| 6 | 8.2, [5, 12] | 5.6, [4, 7] |
| 7 | 8.4, [5, 10] | 6.3, [3, 9] |
| 8 | 9, [5, 14] | 6.4, [3, 10] |
| 9 | 14.3, [13, 18] | 12.4, [10, 16] |
| 10 | 22.7, [16, 29] | 22.3, [18, 28] |

**Supplemental Table 2:** Number of times we measured fractional change in median fixation duration per fish using cluster ablation data (Figure 7D, Supplementary Figure 9D and associated text). Note that the animals with spinal cord ablations are different than the animals with cluster hindbrain ablations

| ANIMAL INDEX | 1 | 2 | 3 | 4 | 5 | 6 | 7 | 8 | 9 | 10 | 11 | 12 | 13 | 14 | 15 | 16 | 17 | 18 | 19 | 20 | 21 | 22 | 23 |
| --- | --- | --- | --- | --- | --- | --- | --- | --- | --- | --- | --- | --- | --- | --- | --- | --- | --- | --- | --- | --- | --- | --- | --- |
| HINDBRAIN ABLATIONS | 1 | 7 | 10 | 5 | 2 | 5 | 10 | 6 | 11 | 6 | 4 | 4 | 5 | 5 | 3 | 4 | 2 | 2 | 8 | 2 | 8 | 9 | 7 |
| SPINAL CORD | 4 | 3 | 3 | 3 | 2 | 3 | 4 | 3 | 1 |  |  |  |  |  |  |  |  |  |  |  |  |  |  |

**Supplemental Table 3:**
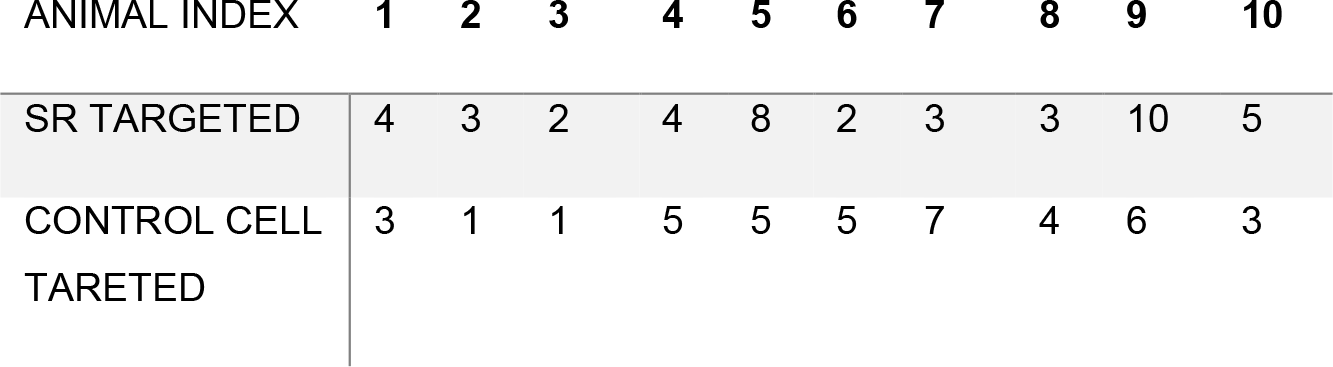
Number of times we measured fractional change in median fixation duration per fish using single-cell ablation data (Figure 7E and associated text). Note that animals with SR-targeted ablations are different than animals with control ablations.

| ANIMAL INDEX | 1 | 2 | 3 | 4 | 5 | 6 | 7 | 8 | 9 | 10 |
| --- | --- | --- | --- | --- | --- | --- | --- | --- | --- | --- |
| SR TARGETED | 4 | 3 | 2 | 4 | 8 | 2 | 3 | 3 | 10 | 5 |
| CONTROL CELL TARETED | 3 | 1 | 1 | 5 | 5 | 5 | 7 | 4 | 6 | 3 |

**Supplemental Figure 1.**
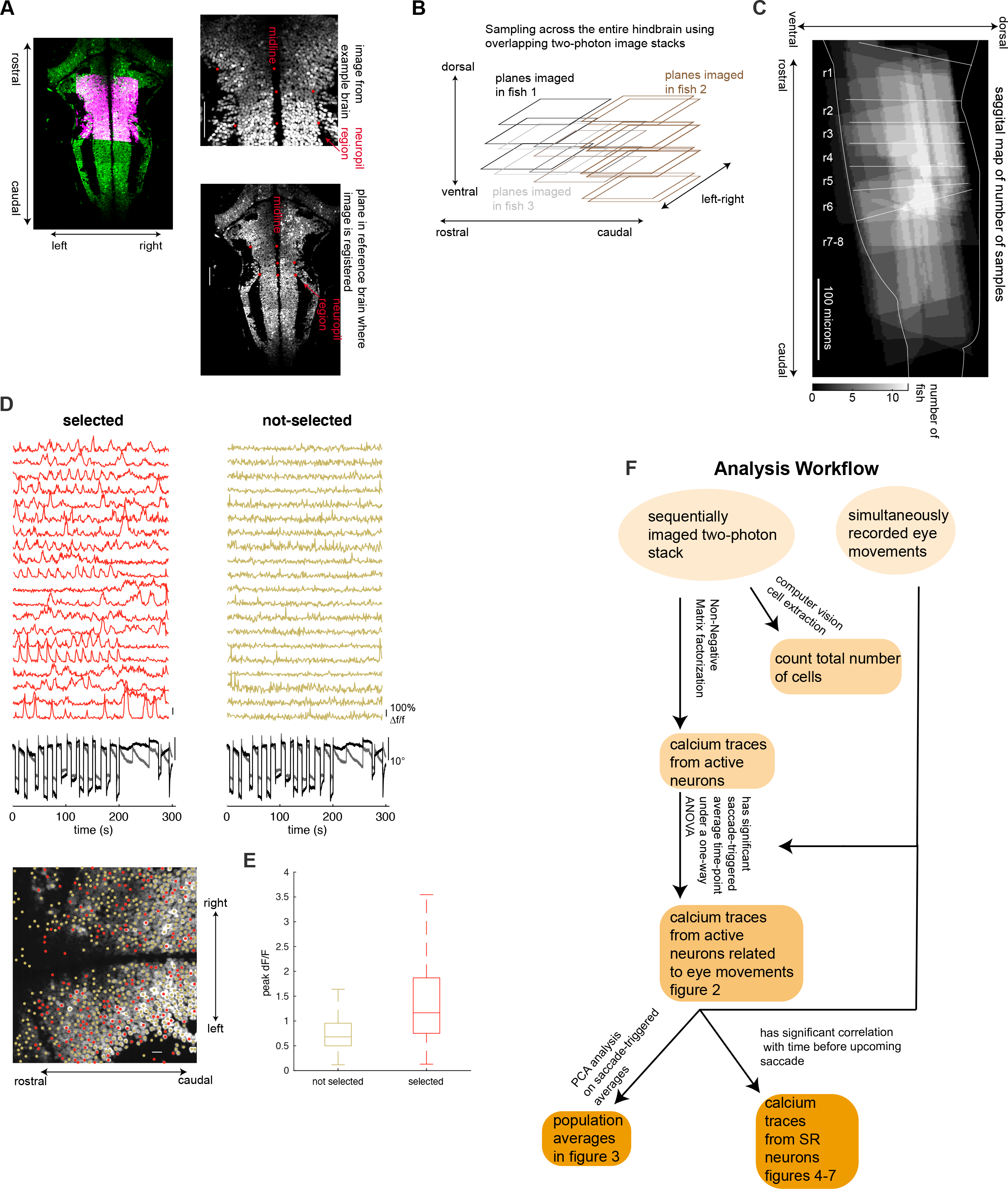
Methods used for imaging and analyzing cells across the hindbrain. (A) Time-averaged calcium images from cells expressing h2b-GCaMP6f demonstrating the procedure used for registration. (left column) Example plane imaged in the rostral hindbrain (pink) after registration overlaid on a corresponding plane in the reference bridge brain (green). (right column) Registration was performed by finding similar points (red dots) between an imaging plane (top) and plane within the reference bridge brain (bottom) and then computing a transformation matrix from these points (see Methods). Similar points were manually found using corresponding features between the example brain and reference brain such as the midline location and neuropil boundaries. This procedure was also performed between the reference bridge brain and Z-Brain atlas. Scale bar length is 50 microns. (B) Cartoon schematic of sampling method used. Each square represents a horizontal imaging plane in the hindbrain. A single imaging plane had a side-length of 185 microns and did not cover the entire rostral-caudal extent of the hindbrain. Overlapping dorsal-ventral stacks of the hindbrain were imaged in multiple fish (different colors) and then registered to a reference brain resulting in hindbrain-wide imaging across the population. (C) Heat map sagittal projection of the number of animals used to sample each location in the hindbrain (20 fish in total). (D) Simultaneously recorded traces of dF/F versus recording time demonstrating that neurons identified by the NMF algorithm as active (red, left column) show greater fluctuations in fluorescence than cells identified as background by the algorithm (yellow, right column). Since the NMF algorithm does not localize background cells we used standard computer vision algorithms to find these neurons based on morphology (cell locations shown by color-coded dots in time-average image of h2b-GCaMP6f at the bottom; scale bar length is 10 microns). Beneath dF/F traces we plot accompanying eye movements. (E) Boxplots of maximum dF/F value across time from a population of cells selected by the NMF algorithm (red) and cells localized by computer vision algorithm and labelled as background by the NMF algorithm (yellow). Central bar shows the median, edges of the box are the 25^th^ and 75^th^ percentiles, whiskers show the range of values except outliers. We rejected the null hypothesis that the average peak dF/F values are equal for the two populations (p<0.01, one-way ANOVA; n=216,717 non-selected cells, n=64,242 selected cells). ANOVA was performed on all cells in the data set not just those registered to the hindbrain as was done throughout the main text. (F) Analysis workflow detailing the steps used to derive cells examined in the manuscript. Videos of two-photon calcium activity were analyzed using the NMF algorithm to identify the locations of neurons that were active during spontaneous activity. Amongst these neurons, we only analyzed cells whose saccade-triggered average (Fig. 2, see Methods) had some variation (tested using a one-way ANOVA). Population-wide patterns of average activity in these cells were characterized using PCA (Fig. 3). Finally, we further selected out cells identified as having pre-saccadic activity which we called SR neurons.

**Supplemental Figure 2.**
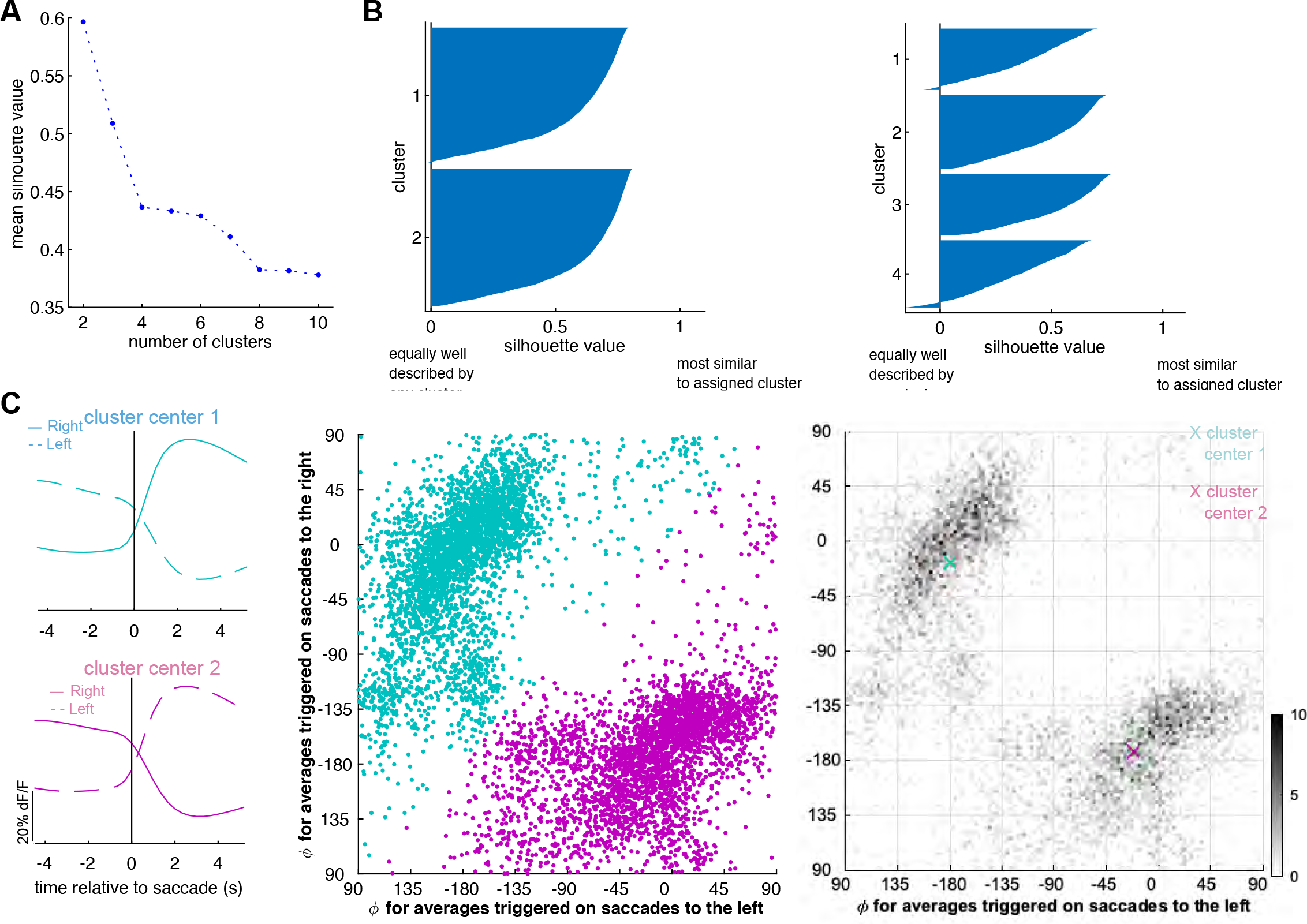
K-Means Analysis of combined left and right saccade-triggered averages. K-means was used to cluster the normalized coefficients found by Principal Component Analysis on saccade-triggered averages (STAs) from eye-movement responsive cells (see Fig. 3Bi,ii and methods section *Principal Component* Analysis for the definition of normalized coefficients). For each cell, we created a 6-dimensional vector by combining the normalized coefficients of the first three components that characterize STAs around saccades to the left and the three that characterize STAs around saccades to the right. To choose the number of clusters, we ran nine K-means analyses, using a different number of clusters on each run (between 2-10) to group the combined coefficients and we used the silhouette value to measure cluster quality. The silhouette value for an individual vector measures how close that vector is to other vectors in its own cluster relative to its distance with vectors in other clusters. The value for the *ith* cell is defined as the minimum average distance from the *ith* vector to all vectors in different clusters than the *ith* vector, *b_i_*, minus the average distance from the *ith* vector to other vectors in the same cluster, a_i_. The silhouette value is normalized by *max*(*a_i_, b_i_*) in order for it to range from +1 to −1 with vectors that are well matched to their cluster having values near +1 and with vectors that are randomly clustered having values near 0. (A) Mean silhouette value versus number of clusters. The largest average silhouette value (best clustering) is achieved using 2 clusters. (B) Silhouette plot showing distribution of silhouette values for all vectors after running K-means using two clusters (left panel) and four clusters (right panel). (C) (left column) STAs around saccades to the left (dashed) and right (solid) corresponding to the cluster means found by K-means analysis using two clusters. The STAs displayed are created by linearly combining the first three components weighted by the coefficients corresponding to the first (top) or second (bottom) cluster center. The center of cluster 1 corresponds to STA pairs that increase in value following saccades to the right and decrease in value following saccades to the left. The center of cluster 2 shows similar activity but with the opposite directional preference. (middle and right columns) For each cell, we plot the angular coefficient, *φ* (see Fig. 3Bii and accompanying methods), characterizing the STA around saccades to the left versus the coefficient characterizing the same cell’s STA around saccades to the right. (middle column) Scatter plot of coefficients with each dot colored according to its assigned cluster after running a K-means analysis on the plot with K=2. Each dot is from a different cell. (right column) Two-dimensional histogram of coefficients as plotted in the middle column (bin size equals 1 degree).

**Supplemental Figure 3.**
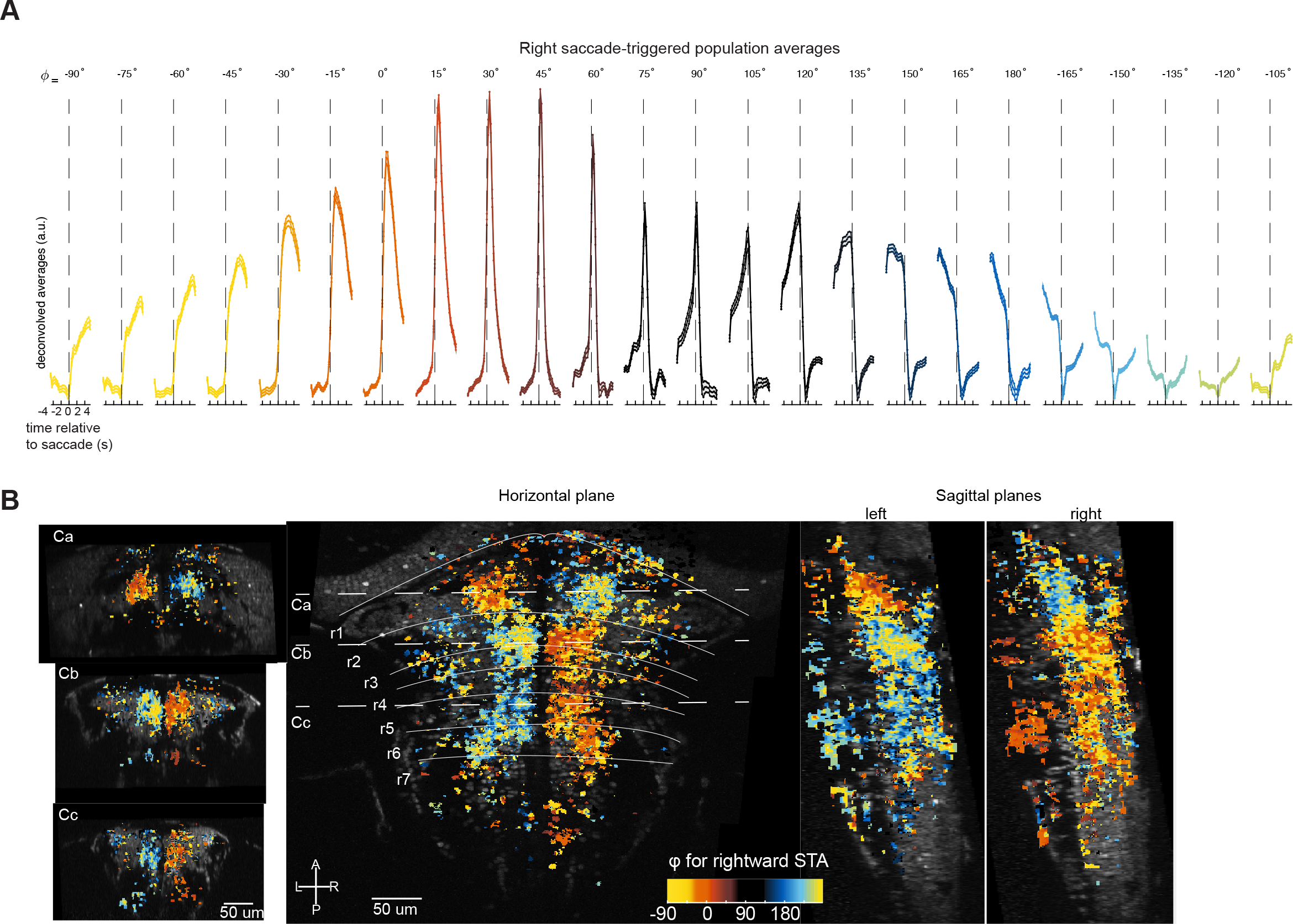
Functional and spatial distribution of eye-movement related activity surrounding saccades to the right. (A) Series of population-averaged STAs after deconvolving with an exponential kernel (2 second time constant). Population averages are comprised of STAs around saccades to the right specified by the value of *ϕ* shown in the top row, (plus or minus 7.5 degrees; see Fig. 3Bi,ii and methods section *Principal Component* Analysis for the definition of *ϕ*). The filled region shows one standard error about the population average. Responses are color coded to enable evaluation of spatial organization in B. (B) Horizontal, sagittal and caudal projections of cell locations color coded by the average value of *ϕ* quantifying STAs around saccades to the right. Color scheme corresponds to the colors shown in A. Coronal sections show average values of *ϕ* for cells within 30 microns of the dashed lines marked C1, C2, and C3 in the horizontal map. Sagittal projections show average values of *ϕ* quantifying STAs around saccades to the right for cells in the left hemisphere or right hemisphere. Number of fish in the averages depends on cell location (see Supplemental Fig. 1C). r, rhombomere; L-R, left-right; A-P, anterior-posterior.

**Supplemental Figure 4.**
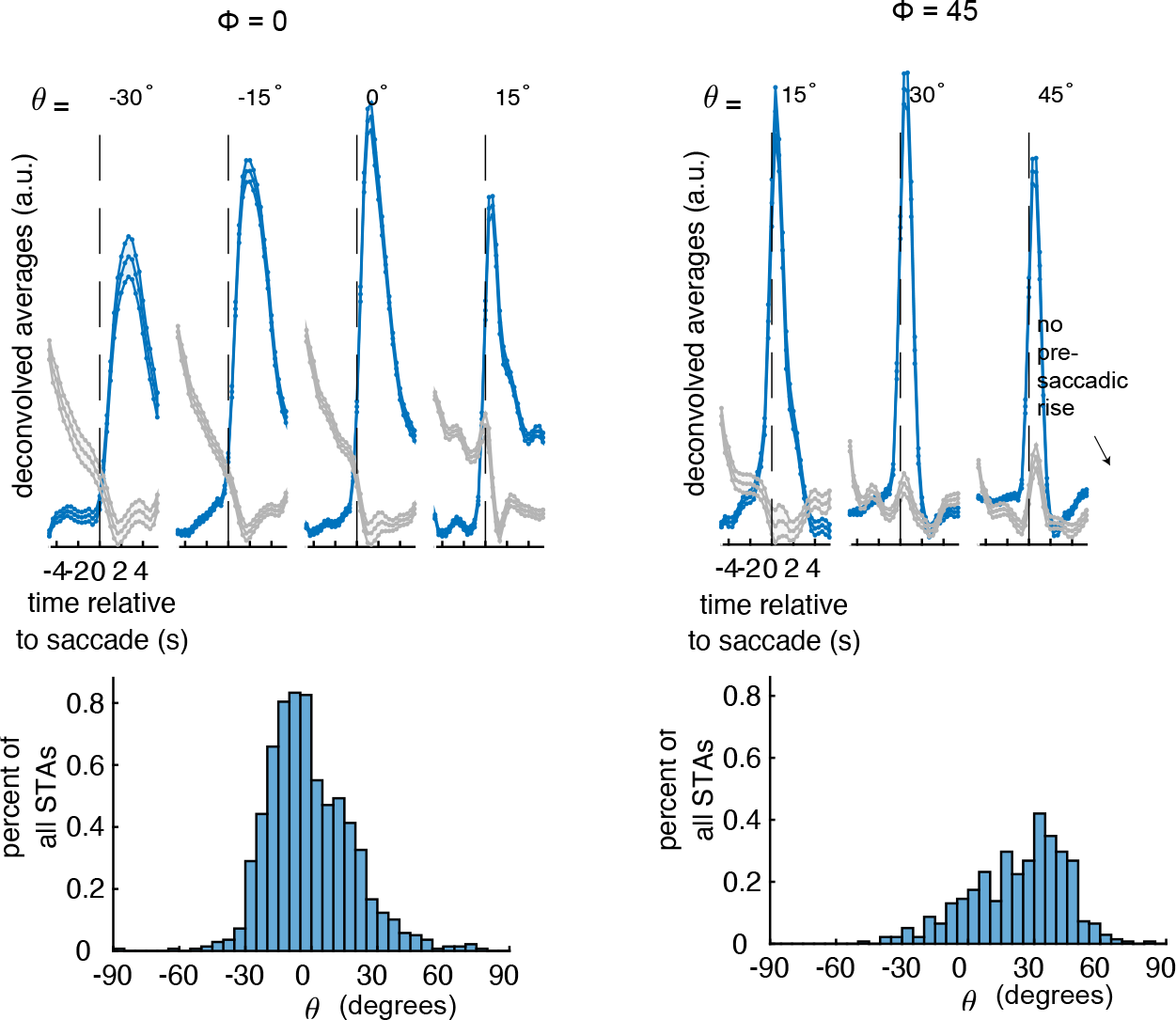
Qualitative features of population average saccade-triggered responses are similar whether responses are binned according to projections along the first three or the first two components provided by principal component analysis. (Top row) Saccade-triggered fluorescence responses averaged across cells after deconvolving with an exponential kernel (2 seconds time constant). We sorted STAs whose value of *φ* was fixed at either 0 ± 7.5 (left column) or 45 ± 7.5 degrees (right column) by their value of *ϑ* (see Fig. 3Bi,ii and methods section *Principal Component* Analysis for the definition of *ϕ* and *ϑ*). The average of STAs with the same value of *ϑ* (within 15 degrees) are shown in blue. We only show population averages for bins of *ϕ*, *ϑ* that contained at least 1% of eye-movement responsive cells. Note that there are no restrictions on whether the STAs averaged together to form the blue trace are around saccades to the left or right. Since single-cells have direction-selective responses (see Supplemental Fig. 2 for further analysis), the two STAs for a given cell will have different temporal profiles (one ON and one OFF) each characterized by different coefficient values and thus a single-cell will only contribute one out of its two STAs to the average. For completeness, we show in gray the population average STA that results when grouping together and then averaging the STAs not included in the blue trace using the same cells. (Bottom row) Histograms of *ϑ* for all saccade-triggered averages with a value of *ϕ* equal to either 0 (left column) or 45 degrees (right column). Here, we only show variations across *ϑ* for two groups of *ϕ* because at other values of *ϕ*, *ϑ* was mostly centered near 0. The population averages shown in Fig. 3C and Supplemental Fig. 3D average together the variation across *ϑ* shown here. In most cases, the averages shown in Fig. 3C and Supplemental Fig. 3D capture the qualitative features of cell populations that have been sub-grouped by *ϑ*.

**Supplemental Figure 5.**
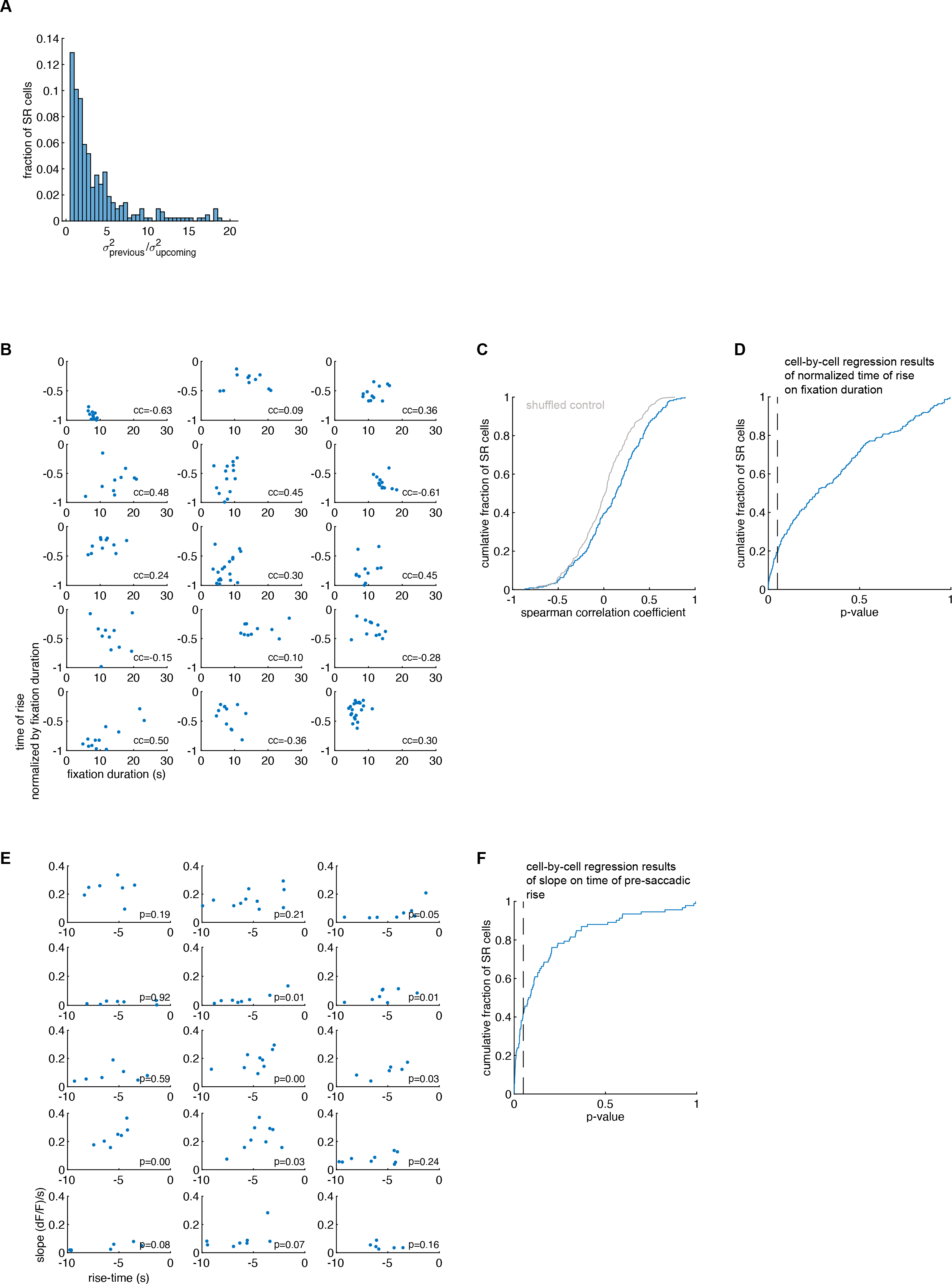
Single-cell analysis of pre-saccadic rise features. (A) For each SR cell we measured the variance across saccades in the time of rise relative to the previous saccade, σ^2^_previous_, and the variance across saccades in the time of rise relative to the upcoming saccade, σ^2^_upcoming_. Histogram across cells of the ratio of variances in time of rise (median ratio = 1.9; n=397 cells). Each cell was required to be recorded along with at least 5 simultaneously recorded saccades. (B) Scatter plots of time of pre-saccadic rise relative to upcoming saccade normalized by fixation duration as a function of fixation duration for 15 example cells. Each subplot corresponds to a different cell. For each cell, we quantified the relationship between the plotted statistics by computing a simple linear regression and Spearman correlation coefficient of normalized rise time on fixation duration. The number at the bottom of each plot reports the correlation coefficient. (C) Cumulative distribution across SR cells of the Spearman correlation coefficients described in B (blue line). Gray line shows the equivalent statistics for controls where we randomly shuffled the normalized time of rise. We rejected the null hypothesis that correlation coefficients across SR cells and shuffled controls come from the same distribution (p<0.001, n=394 cells, Two-Sample KS test). (D) Cumulative distribution across SR cells of the p-values obtained under a t-test with null hypothesis that the regression coefficient described in B is zero. Only 19% percent of single cells had a significant p-value (less than 0.05, dashed line). Amongst significant cells, 83 percent had the same trend (positive slope) as the population. Each cell analyzed in (C) and (D) was required to be recorded along with at least 5 simultaneously recorded saccades and to be recorded when the fixation durations were sufficiently variable (difference between 75^th^ and 25^th^ percentiles of accompanying fixation durations was greater than or equal to 1 second). (E) Scatter plots of slopes of pre-saccadic fluorescence increases as a function of time of pre-saccadic rise for 15 randomly chosen example cells. Each sub-plot corresponds to a different cell (panels in (B) do not necessarily correspond to panels in (E)). For each cell, we quantified the relationship between the plotted statistics using a simple linear regression of slope on time of pre-saccadic rise. The number at the bottom of each plot reports the p-value for the *t* statistic obtained under the null hypothesis that the slope regression coefficient is zero. (F) Cumulative distribution of the p-values described in (E). 29 percent of the cells in the plot have a significant p-value (p<0.05, dashed line) and have the same trend (positive slope) as the population. Each cell was required to be recorded along with at least 5 simultaneously recorded saccades where the slope and time of pre-saccadic rise could be measured and was required to have rise-times that were sufficiently variable for measuring a linear relationship (difference between 75^th^ and 25^th^ percentiles of rise-time was greater than or equal to 1 second; n=326 cells passing criteria).

**Supplemental Figure 6.**
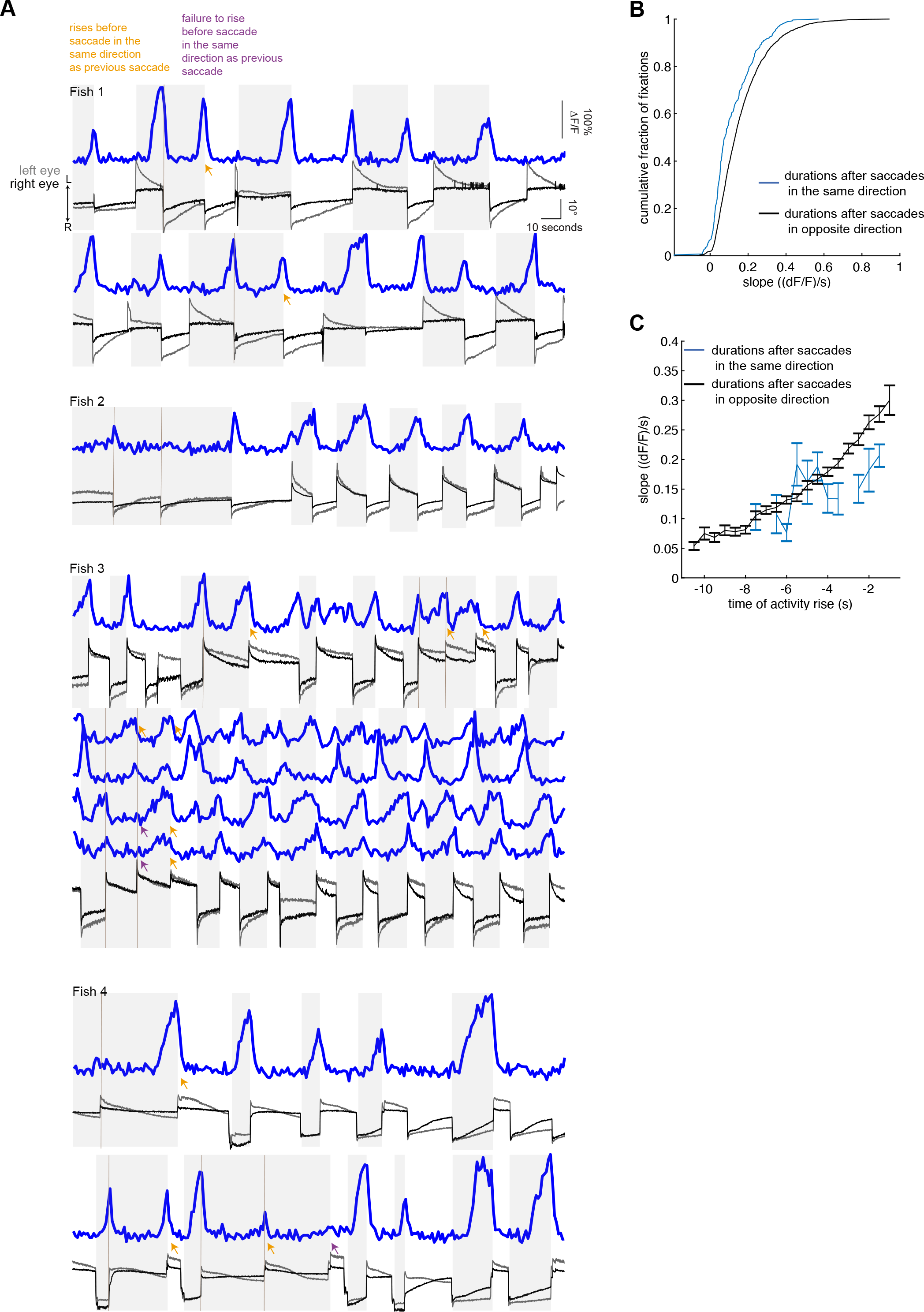
Pre-saccadic rise neuron activity conditioned on saccade patterning. (A) Simultaneously recorded eye position (black and gray traces) and single-cell dF/F responses (blue) from 10 SR cells (found in 4 different fish) recorded when the eyes made at least one saccade in the same direction as the previous saccade. Leftward/rightward directed saccades are made toward the direction labelled L/R. We highlight examples where SR neurons rise before saccades in the same direction as the previous saccade in orange. We also show cases where SR neurons failed to rise before saccades in the same direction as the previous saccade in purple. (B) Cumulative histogram of SR rate of dF/F rise before a saccade in the same direction as the previous saccade (blue) and before a saccade in the opposite direction as the previous saccade (black). (C) Average slope of pre-saccadic fluorescence increases as a function of time of pre-saccadic rise measured with respect to upcoming saccade for saccades made in the same (blue)/opposite(black) direction as the previous saccade. Bin size is 500ms. Bars show mean plus or minus one standard error about the mean.

**Supplemental Figure 7.**
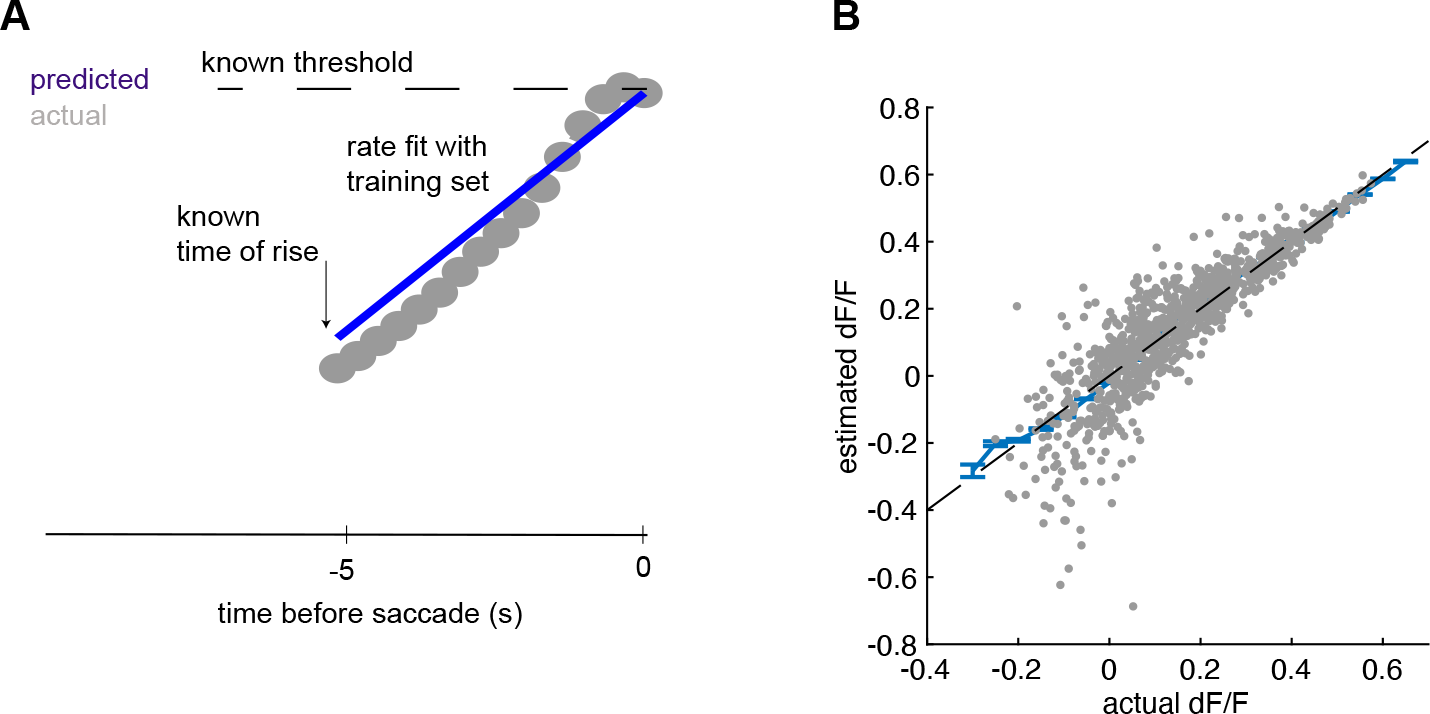
Comparison of SR population average pre-saccadic activity before upcoming saccade with a ramp-to-threshold model. We tested how well SR population average activity fits a model that assumes activity linearly rises from baseline to a threshold value. The model’s rate of rise was fit by constructing a population average from randomly selected SR cells (the training set) and measuring the average rate of rise from baseline before upcoming saccade. The model’s goodness-of-fit was measured by constructing a new population average using SR cells not in the training set and comparing activity with a model that ramps at a rate measured in the training set to a known threshold. (A) Example of population average dF/F from cells in the test set (gray dots) and predicted activity from the ramp-to-threshold model (blue). (B) Quantitative comparison of predicted and actual dF/F activity across 10,000 test sets (using 60% of cells for training and 40% for testing). As seen in (A) the model and actual activity mostly deviate near baseline at values of dF/F near zero (possibly due to non-linear increases in the population as soon as activity begins to rise). However, the model can reproduce population average activity over a large range of dF/F values (r-squared = 0.69). R-squared was measured as one minus the ratio of mean-squared error between predicted and actual dF/F over the variance in dF/F.

**Supplemental Figure 8.**
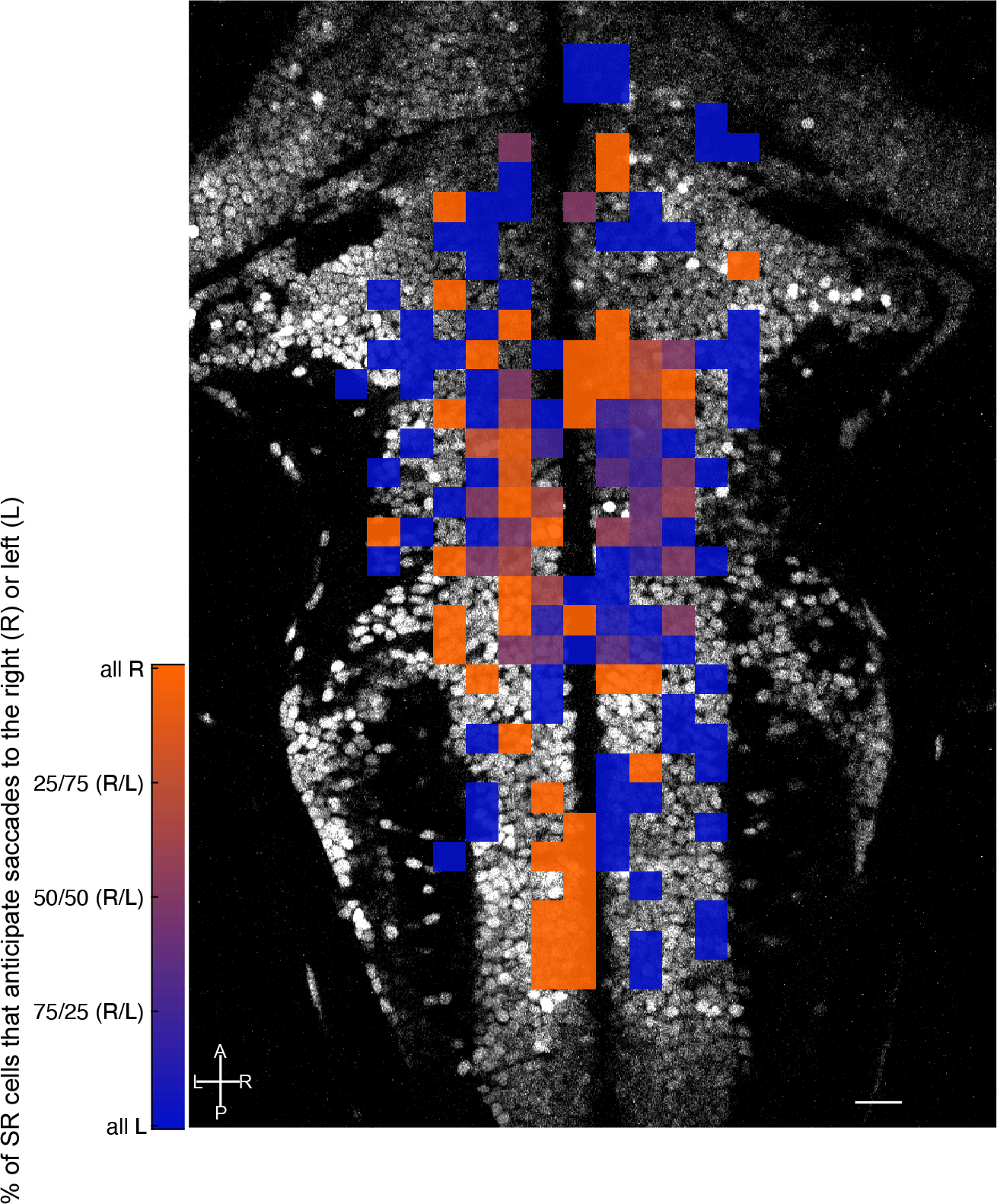
Directional preference of pre-saccadic rise cells. Horizontal projection of directional preference for all pre-saccadic rise cells following alignment with a reference brain. Black and white background image shows one plane from the reference bridge brain for visualization purposes. The color of each 12×12 micron pixel denotes an average of how many cells registered to that square have activity that rises before saccades to the left (blue) or to the right (orange). Squares with violet colors in-between blue and orange indicate that cells with both left and right preference were registered to that location. Scale bar length is 20 microns. A, P, L, R denote anterior, posterior, left and right.

**Supplemental Figure 9.**
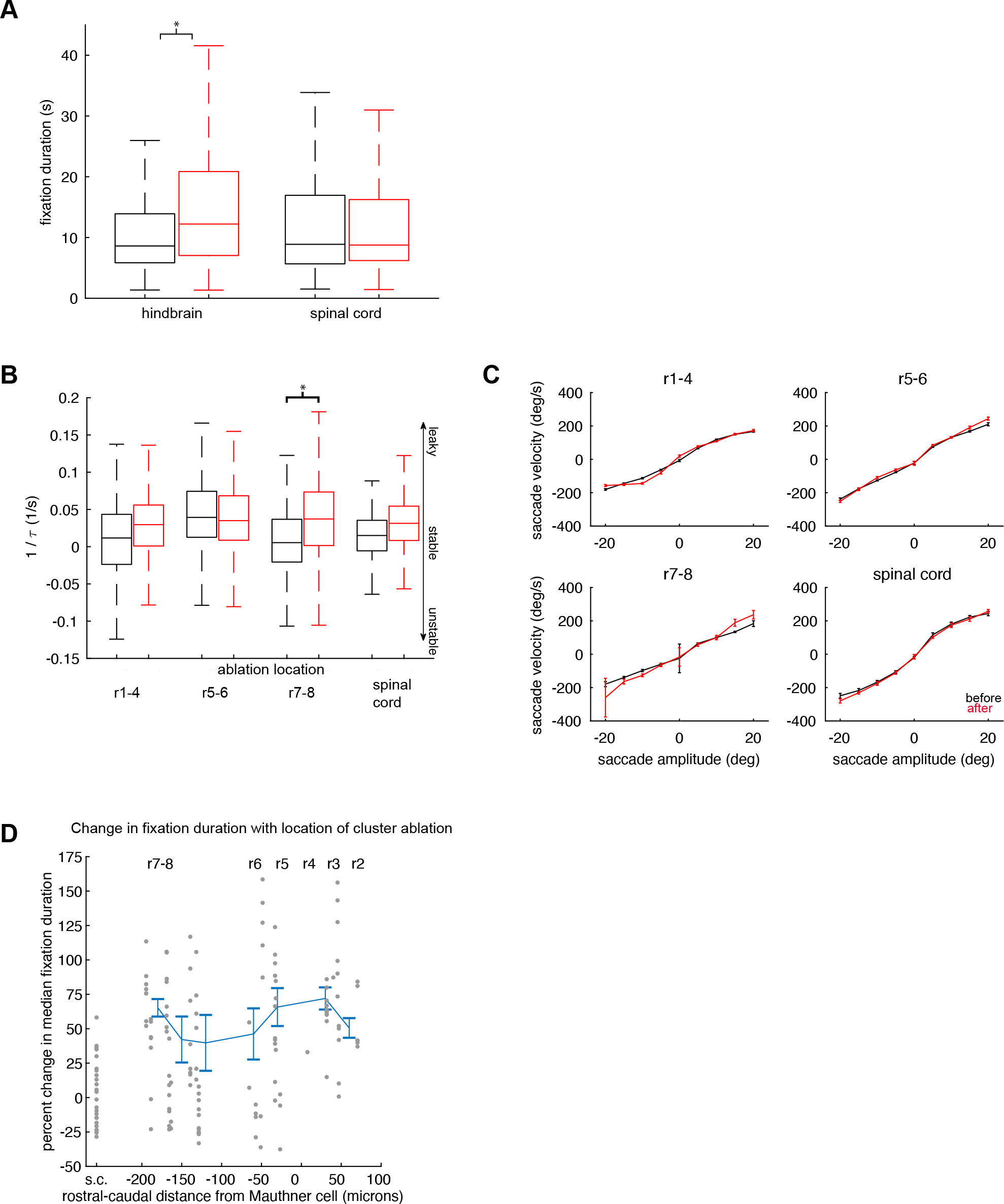
Changes in eye-movement metrics following cluster ablations. (A) Boxplot of fixation durations measured before (gray) and after (red) hindbrain and spinal cord cluster ablations. Left and right eye fixations are combined. Central mark shows the median, edges of the box are the 25^th^ and 75^th^ percentiles, and whiskers show the range of values except outliers. Star shows significant differences. (B) The rate of eye position decay following saccades was measured using an exponential function with time constant tau. Boxplot of inverse tau values before (gray) and after (red) ablations grouped according to ablation location. Large positive values of 1/tau correspond to events where the eyes rapidly returned to nasal resting position. Negative values of 1/tau correspond to events where the eyes continued to move away from nasal resting position after saccade. Central mark shows the median, edges of the box are the 25^th^ and 75^th^ percentiles, whiskers show the range of values except outliers. Stars show significant differences (p<0.001, two-sample t-tests using Bonferroni correction to control for familywise error rate; n=2,053 fixations before and 1,335 fixations after from 6 fish for rhombomeres 1-4 (r1-4), n=1,509 fixations before and 2,038 fixations after from 7 fish for rhombomeres 5-6 (r5-6), n=4,959 fixations before and 2,653 fixations after from 10 fish for rhombomeres 7-8 (r7-8), n=945 fixations before and 1,188 fixations after from 9 fish for spinal cord). (C) Average saccade velocity, grouped according to ablation location, before (gray) and after (red) ablations as a function of saccade amplitude. n=2,060 saccades before and 1,339 saccades after for rhombomeres 1-4, n= 1,510 saccades before and 2,038 saccades after for rhombomeres 5-6, n=4,972 saccades before and n=2,661 saccades after for rhombomeres 7-8, n=949 saccades before and n=1,192 saccades after for spinal cord ablations. Error bars show standard error about the mean. (D) Percent change in median fixation duration following ablation as a function of rostral-caudal position of ablation location. Each gray point shows the percent change in median fixation duration following ablation in a single animal (see Methods). Points at location s.c. are from spinal cord ablated control animals. Average plus standard error about the mean of gray points within 30 micron sized bins (blue line). Bins with less than 10 samples are not plotted.

